# Generalized Maternal-Paternal Genotype Markov Model for Single-Cell Lineages

**DOI:** 10.64898/2026.09.28.755156

**Authors:** Julia Chifman, Adam C. Knapp

## Abstract

Single-cell sequencing provides a rich source of data that can be used to model and estimate the evolution of somatic cells within individuals. Many common models for modeling DNA evolution in phylogenetics are based on continuous-time Markov processes that operate on aligned sites. For haploid DNA, such as found in mitochondria, there are many well established models for these processes but since nuclear DNA is diploid, containing copies of both the maternal and paternal genome, it is natural to develop continuous-time Markov processes adapted to diploid DNA. One possible model is that both the maternal and paternal evolve independently according to haploid models so that, in principle, DNA comes with a *phase*. i.e. a label that tells if the site was derived from the maternal or paternal sequence. However, this information may not be available in practice and only an *unphased* diploid genotype is available. That is, an unphased model groups both of the possible maternal-paternal labels into a single state. This grouping process, called “lumping”, is used by other methods as well and results in a coarser process. Care must be taken so that the coarser process can still be modeled as a continuous-time Markov chain; the lumped process of the original phased Markov chain may or may not be Markovian, depending on restrictions imposed on the generator matrix. Here, we explore the mathematical formulations of phased and unphased Markov processes, as well as a collection of other diploid DNA models from the literature.

## 1 Introduction

Advances in single-cell genome sequencing have led to the development of many probabilistic models that utilize single-cell data to infer cell lineages and the developmental or clonal organization of human tissues and organs, collectively known as human somatic mosaicism (see review articles [1, 2]). While many models were developed with tumor evolution in mind, some of the existing models have been applied to normal hematopoietic stem cell lineages [3] and normal multi-tissue developmental lineages [4]. Single-cell technologies differ in their outputs, ranging from expression profiles to copy-number and genotype characters, all leading to probabilistic models adapted to these states. Models specific to the analysis of single nucleotide variant (SNV) loci can be roughly partitioned into two categories: infinite-sites models (e.g., [5–7]) and finite-sites models (e.g., [8–13]). Some models view each locus in terms of mutation absence or presence, whereas others model genotype states explicitly. The focus of this article is on finite-sites models for diploid genotype evolution. Although many cancers are highly aneuploid, properly classifying diploid models before considering aneuploidy is a first step toward clarifying the basic properties and implications of such models under various assumptions.

The two sources of DNA in a human cell are the mitochondria and the cell nucleus. While mitochondrial DNA is haploid, nuclear DNA is diploid and consists of distinct maternal and paternal genomes. During mitosis, somatic cells pass both mitochondrial DNA and the maternal and paternal nuclear genomes to daughter cells, and in the absence of mitotic recombination, this identity is preserved. Sequence evolution for mitochondrial DNA is typically modeled using a continuous-time Markov process on a collection of aligned sites. Similarly, since both maternal and paternal DNA are passed to daughter cells, it is natural to extend this modeling framework to the maternal and paternal genomes individually. However, because maternal and paternal DNA share a common alignment, these individual processes can be combined to form a *phased* genotype modeled by a continuous-time Markov process of the form *X*(*t*) = (*X*_m_(*t*), *X*_p_(*t*)), with marginal processes *X*_m_(*t*) and *X*_p_(*t*) corresponding to maternal and paternal sequence evolution, respectively.

Phased genotypes are not always produced by sequencing methods and, in some cases, only unphased genotypes can be inferred. That is, we know which alleles are present at a site, but do not know which allele is located on the maternal or paternal chromosome. From a modeling perspective, this suggests building models in which heterogeneous genotypes are grouped together, or lumped, resulting in a coarser process. Such coarser (lumped) processes are also found throughout the literature [8–13], but existing methods do not address the connection between the observable process (unphased genotypes and their various groupings) and the underlying phased process by which maternal and paternal sequences evolve. The significance of this connection cannot be ignored, as the process on the lumped state space may fail to be Markovian, resulting in biased estimates of numerical parameters. Indeed, a recent study reported significant distortions of edge lengths in estimated trees when states were lumped and showed that the resulting process on the smaller state space failed to be Markovian [14]. We further show, in Section 7, that several published continuous-time Markov models cannot be obtained by lumping an underlying independent phased process, but can be obtained by lumping a dependent phased process. In Section 8, we evaluate our independent unphased lumped DNA model against the GT10 model of [12].

## 2 Data Description and Conventions

Genomic data is often stored as a Variant Call Format (VCF) [15] file that records genotypes in terms of variant calls. This essentially provides a list of sites in which at least one taxon was observed to differ from a reference genome. The file also contains other information not discussed in this section, as the focus here is on Markov processes on the genotype space that are suitable for data that exclude non-diploid regions and account only for base substitutions. The VCF file records the observed genotype of each sample at the site in either the form *i* | *j* or *i/j* for diploids, indicating whether the call has been made as phased or unphased, respectively. The characters *i* and *j* are allele values, encoded by non-negative integers. The integer 0 represents the reference allele, and subsequent integers {1, 2, …} represent the first alternate allele, the second alternate allele, and so on. In many VCF files, we will have a mixture of phased and unphased calls, which makes it impossible to assemble a single phased genotype data. Further, reference and alternate genotypes can potentially be complex, with variable numbers of nucleotides. When this is the case, it may be sensible to consider models in which the states are the reference and alternate alleles in their *i/j* forms as specified in the VCF file. However, if the VCF file only consists of single nucleotide polymorphisms (SNPs), we can translate the *i*’s and *j*’s from the VCF file back to nucleotides if desired.

For modeling purposes, we adopt the following notation: phased genotypes will be represented as *i*_m_*j*_p_, where *m* denotes the maternal genome and *p* denotes the paternal genome. For convenience, we have dropped the bar in the middle of the phased genotype. The set of all phased genotypes of size *κ*^2^ will be denoted by

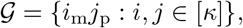

where [*κ*] = {1, 2, …, *κ*}. For unphased genotypes, we use the conventional notation *i/j*, without specifying which allele is maternal or paternal. In particular, the unphased genotype *i/j* represents the set {*i*_m_*j*_p_, *j*_m_*i*_p_}.

### Remark 2.1.

*We can model sequences that have reference, alternate, and second alternate alleles using the set* [*κ*] = {0, 1, 2 *}. The corresponding set of nine phased genotypes is*

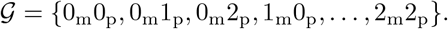

*If one desires to use genotypes, then the elements of* [*κ*] *can be associated with elements of {A, C, G, T} using the REF and ALT fields of the VCF file. This results in 16 ordered pairs*

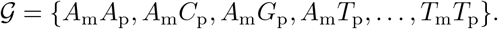

*However, we would like to note that in this case, A*_m_*A*_p_, *for example, could mean* 0_m_0_p_ *or* 1_m_1_p_. *In particular, the distinction between reference and alternates is blurred and site dependent. Which summary of data one should use depends on the questions being asked; however, the Markov process in either case would have the same mathematical structure*.

## 3 Lumpability Overview

As mentioned above, genotype phasing is not always available and so it becomes necessary to group genotypes into a single unphased class *i/j* = {*i*_m_*j*_p_, *j*_m_*i*_p_}. Some models in the literature collapse unphased genotypes even further. For example, Kang et al. [11] combined unphased ternary genotypes into four sets {0/0}, {0/1}, {0/2}, {1/2} and {1/1, 2/2}, represented by 0/0, 0/1, 1/1^*′*^ and 1/1, respectively. The authors refer to set 0/1 as a single mutant and 1/1 as a double mutant, which is biologically meaningful. However, the process with a smaller state space corresponds to a lumped process of the original phased Markov chain, and may or may not be a continuous-time Markov process, depending on restrictions imposed on the generator matrix.

Some notations, definitions, and theorems below concerning lumpability are drawn from Burke and Rosenblatt [16], Kemeny and Snell [17], Ball and Yeo [18]. We do not intend to provide a comprehensive overview of lumpability and the existing literature, and only list the necessary results that we will use for our models.

Let *X*(*t*), *t* ≥ 0, be a homogeneous continuous-time Markov process on the finite state space *S* = {1, 2, …, *n*}. The transition probability matrix is *P* (*t*) = (*p*_*ij*_(*t*)), where under the assumption of time-homogeneity

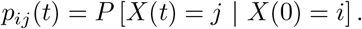

In particular, *P* (*t*) = exp(*Qt*), where *Q* = *P* ^*′*^(0) is the generator matrix of *X*(*t*) with non-diagonal entries *q*_*ij*_ ≥ 0 and diagonal entries *q*_*ii*_ = ∑− _*j*∈*S\{i}*_ *q*_*ij*_.

Let *E* = {*E*_1_, …, *E*_*r*_} be a partition of *S* and let *f* be the function that sends each element of *S* to its partition class. Each of the collapsed states *E*_*i*_ can be viewed as a single state of a new process, 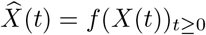, defined over the state space *E*.

### Definition 3.1.

*A continuous-time Markov chain X*(*t*) *is lumpable with respect to E if, for any initial distribution, (1) the lumped process* 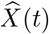 *is a continuous-time Markov chain and (2) the associated transition probabilities do not depend on the choice of initial distribution*.

Some authors differentiate between strong and weak lumpability (e.g. [17, 18]), where strong lumpability is defined as in Definition 3.1 whereas weak lumpability only requires that the lumped process be a continuous-time Markov chain for one initial distribution. Throughout this article, when we refer to lumpability, we will exclusively mean strong lumpability.

Ball and Yeo [18] provide a necessary and sufficient condition for lumpability. In their work, they assume a countable state space and require that continuous-time Markov processes are irreducible and positive recurrent, with some additional assumptions on the generator matrix. The model that we introduce and describe in Section 4 is a finite-state, irreducible, continuous-time Markov process, and hence positive recurrent and conditions on the generator matrix hold trivially. Thus, the results of Ball and Yeo [18] apply in our setting.

Let *Q*_*ij*_ be a submatrix of *Q* corresponding to the classes *E*_*i*_ and *E*_*j*_. The entries of *Q*_*ij*_ are given by *q*_*kn*_, with *k* ∈ *E*_*i*_ and *n* ∈ *E*_*j*_. For example, if *S* = {1, 2, 3, 4, 5} is partitioned by *E*_1_ = {1, 2} and *E*_2_ = {3, 4, 5}, then the generator matrix *Q* is partitioned as follows:

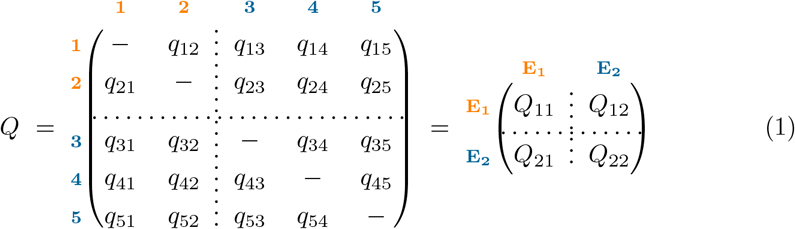

Here and throughout, for rate matrices, we use the convention that the diagonal entries are represented by “–”, as they are simply the negative sums of the non-diagonal entries in the same row.

### Theorem 3.2.

*Ball and Yeo [18, Theorem 2*.*4] The process X*(*t*) *is strongly lumpable if and only if, for all i, j, there exists λ*_*ij*_ ∈ ℝ *such that*

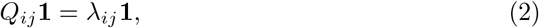

*where* **1** *is a column vector of ones of appropriate dimension*.

This means that each row of *Q*_*ij*_ must sum to the same value, that is

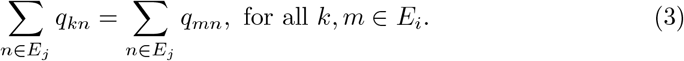

Relation 3 also holds for the transition probability matrix *P* (*t*) = exp(*Qt*). This means that a necessary and sufficient condition for *X*(*t*) to be lumpable with respect to *E* is that 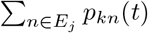 has the same value for all *k* ∈ *E*_*i*_, for every pair *E*_*i*_, *E*_*j*_ ∈ *E*. (See [16] for continuous-time, and [17] for discrete-time.)

We single out Theorem 3.2 as it intuitively leads to a characterization of lumpability in terms of matrices, which we utilize throughout this article. Equation 2 can be expressed in matrix form as follows. Define an *n × r* matrix *V* = (*v*_*ij*_) by

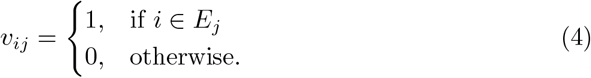

Let 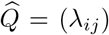 be an *r × r* matrix with 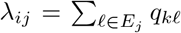 for *k* ∈ *E*_*i*_, cf. Equation 2. The *λ*_*ij*_ represent instantaneous rates of change from state *k* into a partition *E*_*j*_ and have the same value for every *k* ∈ *E*_*i*_ under lumpability. Since *Q*_*ii*_**1** = − ∑ _*i*≠*j*_ *Q*_*ij*_**1**, this ensures that *λ*_*ii*_ is the negative sum of the non-diagonal entries in the same row. Then the full matrix form of Equation 2 is 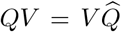. Using our running example and conditions of Theorem 3.2 we get:

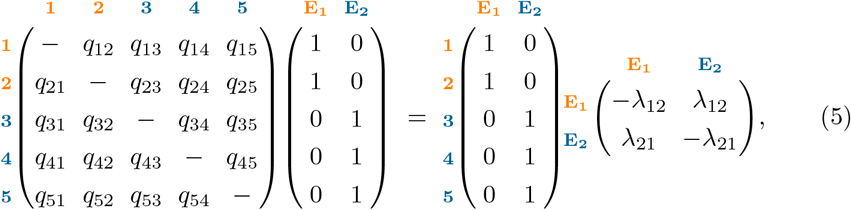

where *λ*_12_ = *q*_13_ +*q*_14_ +*q*_15_ = *q*_23_ +*q*_24_ +*q*_25_ and *λ*_21_ = *q*_31_ +*q*_32_ = *q*_41_ +*q*_42_ = *q*_51_ +*q*_52_.

### Remark 3.3.

*For any r × n matrix U such that UV* = *I*_*r*_, *we have* 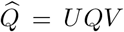. *Thus, if the process X*(*t*) *is lumpable, then the matrix* 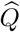 *is the generator matrix of the lumped process* 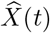.

While the precise definition of *U* varies in the literature, the result is algebraic and does not affect the lumpability conditions of Theorem 3.2, and leads to the same lumped 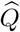. Some authors require that *U* is constructed using an initial distribution with all entries positive [16], which is equivalent to requiring that entries in row *i* corresponding to the states in *E*_*i*_ are non-zero and sum to one, while all other entries are zero [17]. In principle, the matrix *U* can be any matrix where the entries in row *i* corresponding to the states in *E*_*i*_ sum to one, and all other entries are zero. This means that entries within the same partition can be zero as long as their sum equals one.

Throughout this article, we use a *U* equal to the Moore-Penrose pseudoinverse:

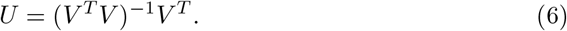

Row *i* of *U* has equal weight for all states within each partition *E*_*i*_ and zeros otherwise. This choice of *U* simplifies many of the arguments that follow.

### Remark 3.4.

*From the discussion above, for any r × n matrix U such that* 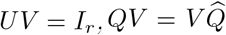 *if and only if V UQV* = *QV*. *Combining this with Theorem 3*.*2, see that the process X*(*t*) *is lumpable if and only if*

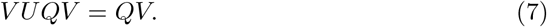

*Note that* 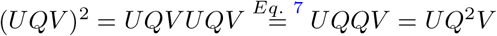. *So, by induction, we see that*

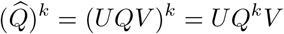

*for any non-negative integer k. Then*,

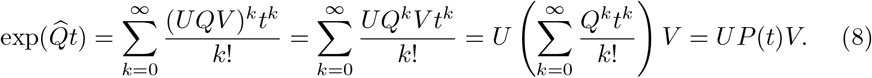

*Also note that* 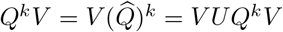, *leading to V UP* (*t*)*V* = *P* (*t*)*V*. *Let*

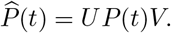

*Then* 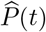 *is the transition probability matrix for the lumped process* 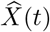.

Recall that, if the process *X*(*t*) on the finite state space *S* is irreducible, then there is a unique stationary distribution *π* = (*π*_1_, …, *π*_*n*_), which is also a limiting distribution. Then lim_*t*→∞_ *P* (*t*) = **1***π*, where **1** is the column vector of all ones in ℝ^*n*^. Thus, **1***π* is a limiting matrix whose rows are all equal to *π*. Also, since *π* is stationary, *πQ* = **0**. Now suppose that *X*(*t*) is lumpable with respect to *E*. Then 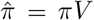 is a stationary distribution for the lumped process 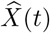. This follows from a simple computation:

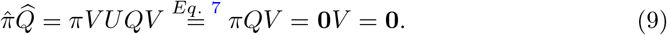

Since both *U* and *V* do not depend on time, 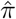 is also a limiting distribution for the lumped process. Thus,

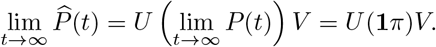

The discussion below Remark 3.3 shows that *U* **1** is a column vector of all ones in ℝ^*r*^, where *r* is the number of classes in *E*. Thus, the rows of *U* **1***πV* are given by *πV*.

## 4 Model Description

The models mentioned in Section 1 incorporate a finite-sites continuous-time Markov model of sequence evolution within either a maximum likelihood or Bayesian framework. Regardless of the framework, many methods require aligned sequences and, in this case, aligned genotypes. This results in a data matrix whose rows represent individual cells and whose columns represent genotypes. Each column is referred to as a *site*, and one common assumption is that sites evolve independently. This allows the likelihood to be written as a product of the conditional probabilities of the data at each site, with a common formula dependent on a given phylogenetic tree. Evolution in different lineages, that is, along different branches of the phylogenetic tree, is also assumed to be independent. This, in turn, allows us to focus on modeling genotype evolution over a single branch, as one sequence evolves into another, and at an individual site.

The genotype Markov model that we describe in this section does not account for indels and models only substitutions. There are attempts in the cancer phylogenetics community to use deletions (“–”) as a possible state. For example, in Kang et al. [13], these unphased genotypes are written in the form 0/− or 1/−. However, while a deletion can occur at a single nucleotide base pair, it can also span many bases. Therefore, it is not immediately clear how the assumption of site independence affects a single deletion event that spans many bases. For example, if a deletion spanning 100 bases is treated as 100 independent site-level deletion states, the effect on parameter inference is unclear. This is an important question to address and is part of our future work, but in this article we consider only substitutions.

Let *X*(*t*) = (*X*_m_(*t*), *X*_p_(*t*)) be a continuous-time Markov process on the phased diploid genotype state space *G* of finite size *κ*^2^, describing genotype-level sequence evolution along a single phylogenetic-tree branch. The component processes *X*_m_(*t*) and *X*_p_(*t*) correspond to marginalized maternal and paternal sequence evolution, respectively, and are not necessarily Markovian. Let 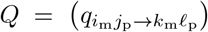 be the *κ*^2^ *× κ*^2^ generator matrix of *X*(*t*), describing the instantaneous rates of change from genotype *i*_m_*j*_p_ to genotype *k*_m_ℓ_p_. We assume that the evolutionary process is time-homogeneous and that *X*(*t*) is irreducible. Hence, there is a unique stationary distribution, which is also the limiting distribution. Other restrictions may be imposed as needed. Branches of phylogenetic trees are typically scaled so that branch lengths correspond to the expected number of mutations per site. Scaling is discussed further in Section 4.6.

Given the description above, it is tempting to just rename genotypes with labels between 1 and *κ*^2^ and use a general *κ*^2^-state Markov model on phylogenetic trees. However, this approach overlooks certain subtleties, which we highlight in this article. Several questions naturally arise: what restrictions are imposed on the generator matrix *Q* if (i) the marginal processes *X*_m_(*t*) and *X*_p_(*t*) are Markovian and independent; (ii) the marginal processes are Markovian but dependent; and (iii) the marginal processes are not Markovian and only *X*(*t*) is a continuous-time Markov process? While all three questions are important, this article focuses primarily on the first. Section 6 provides illustrative examples of cases (ii) and (iii). A detailed exploration of models with dependent or non-Markovian marginals is left for future investigation.

### 4.1 Independent Phased Maternal and Paternal Process

The transition probabilities of the homogeneous continuous-time Markov process *X*(*t*) = (*X*_m_(*t*), *X*_p_(*t*)) are

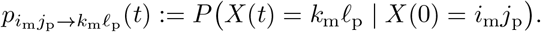

Since *X*(*t*) consists of maternal and paternal components, the transition probability can equivalently be viewed as a joint probability for *X*_m_(*t*) and *X*_p_(*t*). If the maternal and paternal components are independent, then the phased transition probabilities decompose as

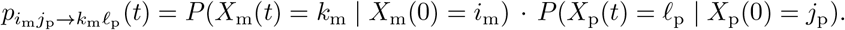

In other words, we can write 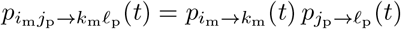, that is, as a product of the transition probabilities of the component processes, which are themselves homogeneous continuous-time Markov processes. Using the Kronecker product, this relationship can be written compactly as

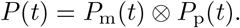

Differentiating at *t* = 0 yields

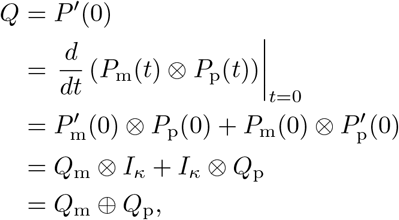

where the third equality follows from the product rule for the Kronecker product and the last from the definition of the Kronecker sum.

For example, consider *κ* = 3 with states {0, 1, 2}. The genotype state space is *G* = {0_m_0_p_, 0_m_1_p_, …, 2_m_2_p_}. Graphically, *Q*_m_ ⊕ *Q*_p_ corresponds to the Cartesian product of the two graphs shown in Figure 1A–B. For this example, the generator matrix of the independent phased process is

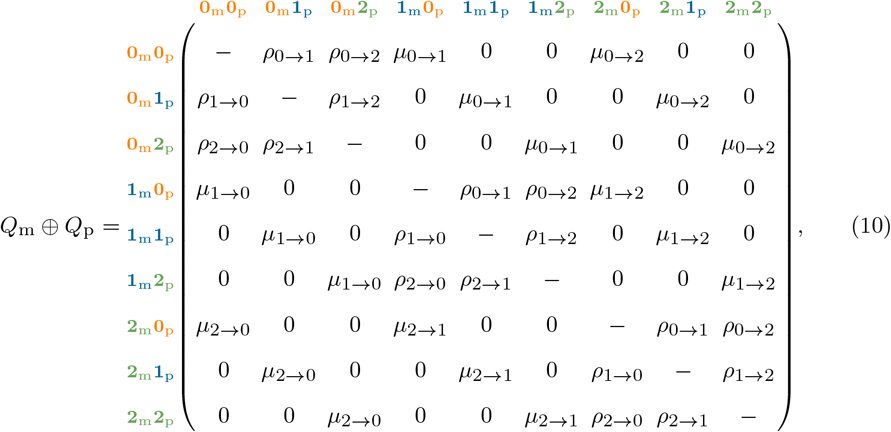

where *Q*_m_ = (*µ*_*i*→*j*_) and *Q*_p_ = (*ρ*_*i*→*j*_) are both 3 *×* 3 generator matrices for maternal and paternal processes, respectively.

**Fig. 1:**
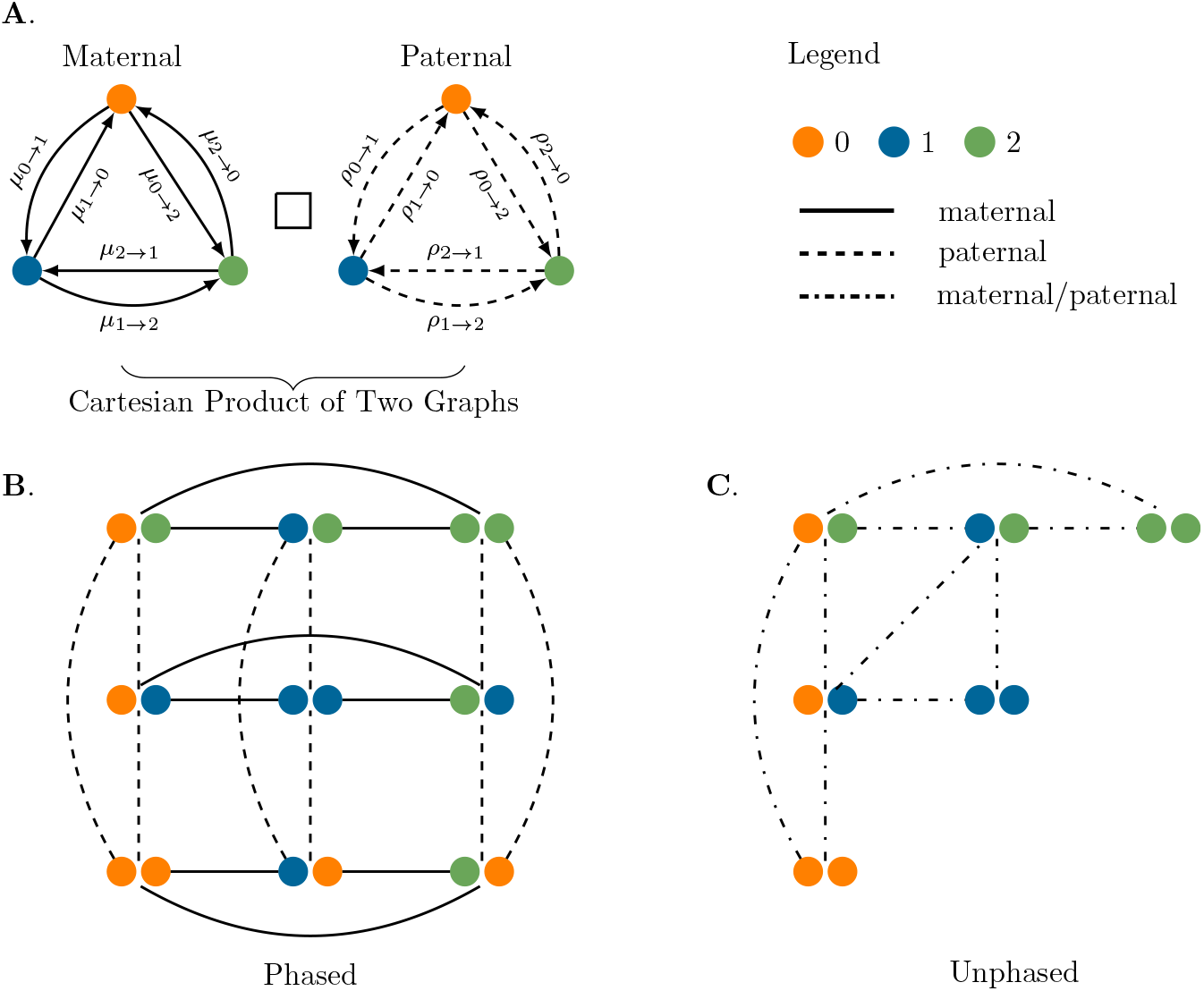
Independent phased/unphased Markov genotype process. (A) Rate diagram for the maternal and paternal processes. (B) The graph representation of the phased model for the joint independent processes is the cartesian product, in the category of graphs, of the graphs for the maternal and paternal processes. Transitions in the maternal sequence are drawn with solid lines and transitions in the paternal sequence are drawn with dashed lines. (C) The graph representation of the unphased model fuses states that are in the same partition. Each transition in the phased model leads to a transition in the unphased model where the rates are derived from the phased model. See e.g. Equation 31 in Section 4.2. This means that each transition, drawn with dot-dash lines, can be the result of either a change in the maternal or paternal sequence.

#### Definition 4.1.

*Let X*(*t*) = (*X*_m_(*t*), *X*_p_(*t*)) *be a continuous-time Markov process on the finite state space G. Let* 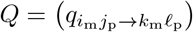 *be the κ*^2^ *× κ*^2^ *generator matrix of X*(*t*) *describing the instantaneous rates of change from genotype i*_m_*j*_p_ *to genotype k*_m_ℓ_p_.

*The process is said to be an* ***independent phased maternal-paternal process*** *if*

1. 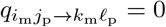 *whenever i*_m_ ≠ *k*_m_ *and j*_p_ ≠ ℓ_p_,
2. *for all j*_p_ *and* 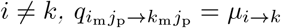 *for a fixed constant µ*_*i*→*k*_, *and*
3. *for all i*_m_ *and* 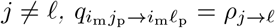 *for a fixed constant ρ*_*j*→ℓ_.

*In particular, the component processes X*_m_(*t*) *and X*_p_(*t*) *are independent continuous-time Markov processes with generator matrices given by Q*_m_ = (*µ*_*i*→*j*_) *and Q*_p_ = (*ρ*_*i*→*j*_), *respectively. The generator matrix of X*(*t*) *is then Q*_m_ ⊕ *Q*_p_.

A common assumption in the literature (e.g., [11, 12]) is that the genotype *i*_m_*j*_p_ does not simultaneously change both component states in an infinitesimal amount of time. This assumption corresponds to the first of the rate conditions in Definition 4.1, that is 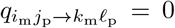 whenever both *i*_m_ ≠ *k*_m_ and *j*_p_ ≠ ℓ_p_. For example, in the phased diploid binary state space, {00, 01, 10, 11}, the rates *q*_00→11_, *q*_01→10_, *q*_10→01_, and *q*_11→00_ are all zero.

A version of the above discussion appears in Ball and Yeo [18, Theorem 3.2], which shows that, assuming the rates of simultaneous changes are fixed at zero, *X*_m_(*t*) and *X*_p_(*t*) are both continuous-time Markov processes if and only if they are independent. We would like to emphasize a subtlety here; it is possible for the marginal processes to be Markovian yet dependent. In such a case, Ball and Yeo [18, Theorem 3.2] implies that the rate of simultaneous maternal-paternal genotype change must be positive. We present some processes with dependent maternal-paternal components in Section 6.

#### 4.1.1 A Few Words About Marginal Processes

Marginal processes are linked to lumpability. In this section, we demonstrate that *Q*_m_ ⊕ *Q*_p_ is lumpable under specific partitions of the genotype space corresponding to the marginalization. In this partition, the lumped process is precisely the maternal (paternal) process, with the generator matrix *Q*_m_ (*Q*_p_). We provide the procedure for the maternal process; the paternal process follows similarly.

Partition the state space *G* of the phased process *X*(*t*) into *κ* disjoint sets

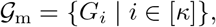

where

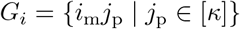

is the set of all phased genotypes with maternal genotype *i*. Thus, each *G*_*i*_ corresponds to state *i* of the maternal marginal process *X*_m_(*t*).

Let **1**_*κ*_ = (1, 1, …, 1) be a length *κ* row vector of all 1’s. Using the lexicographic ordering of phased states, Equations 4 and 6 give

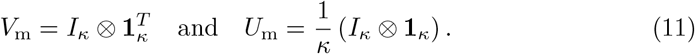

The subscript m is here to emphasize that we are defining matrices for the maternal process. For example, if *κ* = 3 then

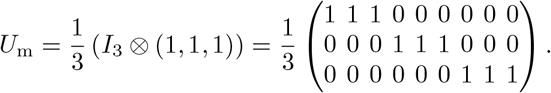

Recall that *U*_m_ is the left inverse of *V*_m_ and so *U*_m_*V*_m_ = *I*_*κ*_. Additionally, we need the product in the other order and can compute

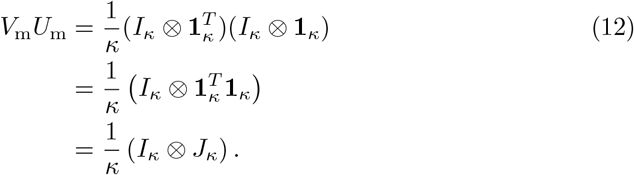

In the equation above, *J*_*κ*_ is a matrix of size *κ × κ* consisting of all 1’s.

To show that *Q*_m_ ⊕ *Q*_p_ is lumpable with respect to partition *G*_m_, we use Remark 3.4.

First, we note that

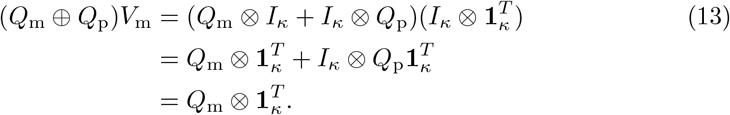

Note that 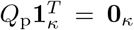, since multiplying by 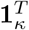 sums the rows of *Q*_p_. Now, using Equations 12 and 13 we get

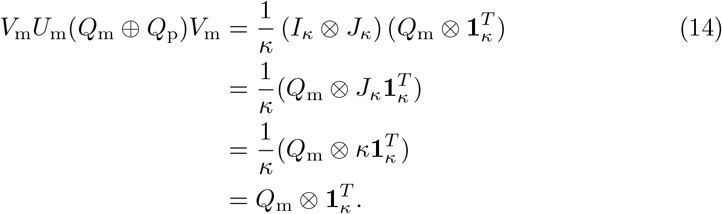

We conclude that *V*_m_*U*_m_(*Q*_m_ ⊕ *Q*_p_)*V*_m_ = (*Q*_m_ ⊕ *Q*_p_)*V*_m_. In particular, *Q*_m_ ⊕ *Q*_p_ is lumpable with respect to partition *G*_m_ and the generator matrix for the lumped maternal marginal process *X*_m_(*t*) is given by

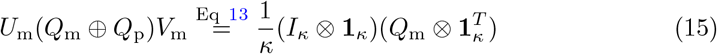

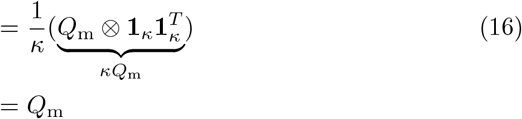

By Remark 3.4, the transition probability matrix for the marginal maternal process is

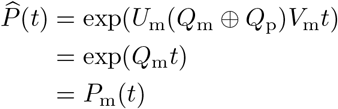

The paternal case follows similarly with *V*_p_ and *U*_p_ defined as

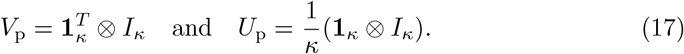

### 4.2 Independent Unphased Maternal and Paternal Process

As many genotype datasets are at least partially unphased, it is important to construct a rate matrix for unphased genotypes. While it is possible to construct *ad hoc* rate matrices that satisfy various natural requirements [11, 12], rate matrices derived from an underlying phased genotype model should be a better match to biological reality. To achieve this, we need to know what restrictions must be imposed on the rate matrix of the phased independent process to ensure that the lumped process is Markovian.

Using notation from Section 2, let *i/i* = {*i*_m_*i*_p_} and *i/j* = {*i*_m_*j*_p_, *j*_m_*i*_p_} _*i*≠*j*_. The lumped state space for the unphased process is then the collection

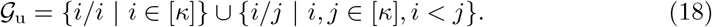

The size of *G*_u_ is 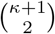 in total with *κ* of the form *i/i* and 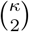 of the form *i/j*. To analyze the unphased process more conveniently, it will be useful to choose an alternative ordering of the *phased* state space which is adapted to the unphased lumping. Reordering the elements of the phased state space permutes the rows of *V*_u_, the columns of *U*_u_, and both the rows and columns of *Q*_m_ ⊕ *Q*_p_. Throughout, we fix the ordering of the unphased states to the one used in the definition (18) of *G*_u_, thereby fixing the columns of *V*_u_, the rows of *U*_u_, and both the rows and columns of the unphased rate matrix 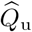.

When we need to distinguish matrices written using the lexicographic ordering of the phased state space, we will use a lex superscript. Otherwise, we order the phased state space as follows. First, the states *i*_m_*i*_p_ are listed in lexicographic order. The remaining states are ordered according to the ordering of the unphased states: for each *i < j*, the pair *i*_m_*j*_p_ and *j*_m_*i*_p_ appears consecutively, with the pairs ordered lexicographically by (*i, j*). These ordering choices lead to a useful block form for *V*_u_ shown in Equation 24. For example, if *κ* = 3 with {0, 1, 2} then the new ordering of phased states is

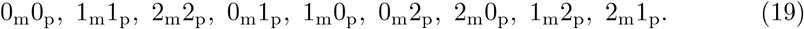

Let Σ be the permutation matrix which realizes the change to the new ordering of the phased state space. In the new ordering, the independent phased process is generated by *Q* = Σ(*Q*_m_ ⊕ *Q*_p_)Σ^*T*^. The matrix *Q* is now in the block form

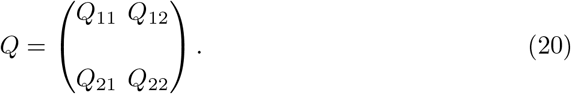

1. The submatrix *Q*_11_ is a diagonal submatrix of size *κ × κ*. The diagonal entry in each row is the negative sum of the corresponding off-diagonal entries of *Q*; these are the rates from the phased genotype of the form *i*_m_*i*_p_ to itself.
2. The submatrix *Q*_12_ has size 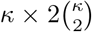. It describes the rates from a genotype of the form *i*_m_*i*_p_ to genotype *k*_m_ℓ_p_, *k* ≠ℓ. The column of *Q*_12_ labeled by *k*_m_ℓ_p_ will only have two non-zero entries in rows *k*_m_*k*_p_ and ℓ_m_ℓ_p_ with rates *ρ*_*k*→ℓ_ and *µ*_ℓ→*k*_, respectively, where *ρ* stands for paternal and *µ* for maternal rates.
3. Similarly, the 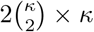 submatrix *Q*_21_ records the rates from *k*_m_ℓ_p_ to *i*_m_*i*_p_, *k* ≠ ℓ, with each row labeled by *k*_m_ℓ_p_ only having two non-zero entries in column *k*_m_*k*_p_ with the rate *ρ*_ℓ→*k*_, and in column ℓ_m_ℓ_p_ with the rate *µ*_*k*→ℓ_.
4. The submatrix *Q*_22_ has size 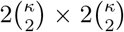 and contains rates between genotypes *i*_m_*j*_p_ and *k*_m_ℓ_p_, with *i* ≠ *j* and *k* ≠ ℓ.

Continuing our running example on 9 states, the *Q* matrix in the block form is:

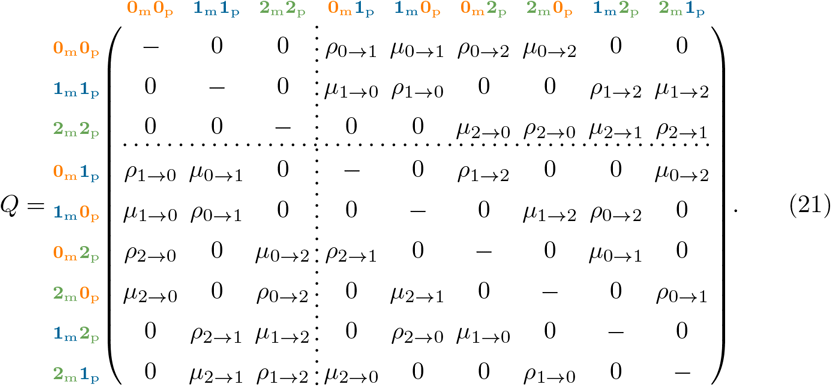

The matrices *V*_u_ and *U*_u_ can be written in block form in terms of **0**_*m×n*_, a matrix of all zeros, and **1**_2_ = (1, 1).

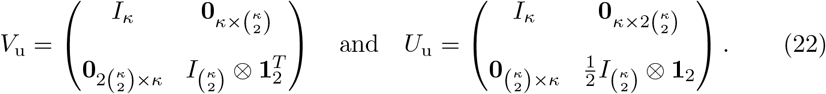

For our running example, *V*_u_ is

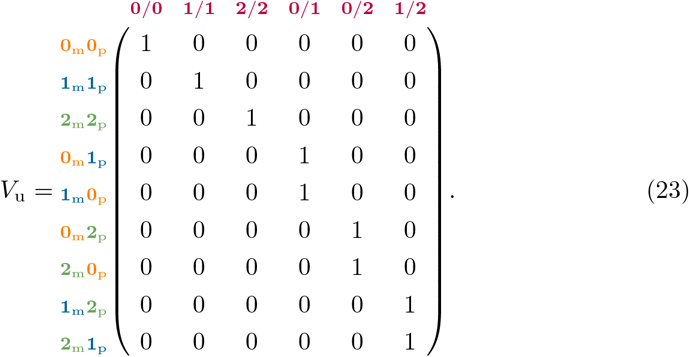

For the phased process to be lumpable with respect to partition *G*_u_, we need *V*_u_*U*_u_*QV*_u_ = *QV*_u_. We check this now.

From now on, except for the identity matrix, we will not specify the size of each block matrix to keep the notation cleaner. Note that *V*_u_*U*_u_ is a square block-diagonal matrix of the form

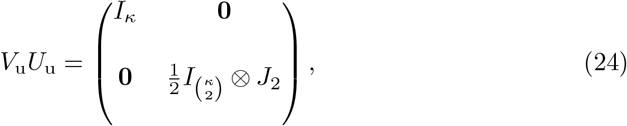

where *J*_2_ is a 2 *×* 2 all-ones matrix. Then *V*_u_*U*_u_*QV*_u_ and *QV*_u_ are

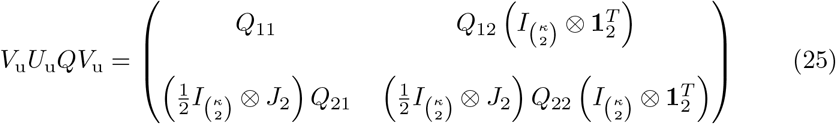

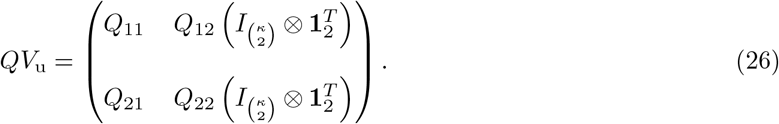

The first row of Equations 25 and 26 are identical. Thus, we focus only on the submatrices of the second row.

#### The (2, 1)-block

The submatrix *Q*_21_ can be viewed as a row-block matrix with 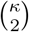 blocks of size 2 *× κ*. The rows of each block are labeled by the states in *i/j* = {*i*_m_*j*_p_, *j*_m_*i*_p_}, *i < j*, and columns are labeled by *k*_m_*k*_p_. There are only two columns out of *κ* columns in each row-block that are non-zero having the entries as shown below:

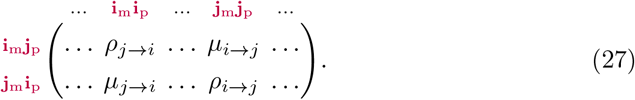

Now, 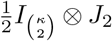 is a block-diagonal matrix with diagonal blocks of the form

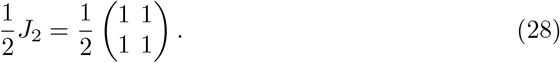

For the following equality to hold,

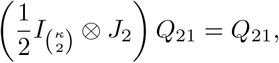

we must have that,

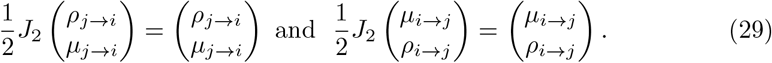

This implies that *µ*_*i*→*j*_ = *ρ*_*i*→*j*_ for all *i, j* ∈ [*κ*], *i* ≠ *j*. That is, *Q*_m_ = *Q*_p_.

#### The (2, 2)-block

Note that *Q*_22_ submatrix is a block-matrix with 2 *×* 2 blocks. Consider a row of *Q*_22_ labeled by the states in *i/j* = {*i*_m_*j*_p_, *j*_m_*i*_p_ *}, i, j* ∈ [*κ*] with *i < j*. Depending on the choice of *i* and *j* the possible structures of these blocks are:

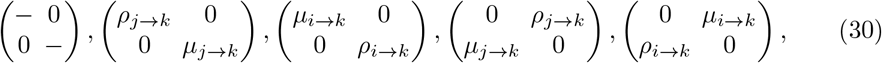

The first block corresponds to *i/j* → *i/j* transitions, while the others correspond to either *i/j* → *k/j* or *i/j* → *i/k* transitions, with the particular block structure depending on the relative ordering of *i, j*, and *k*. The matrix 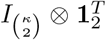 is a block diagonal with diagonal entries of the form 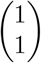. Upon multiplication of *Q*_22_ by 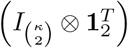, each of the blocks in Equation 30 becomes the 2 *×* 1 blocks.

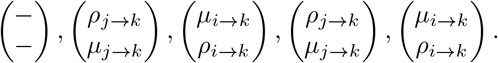

From our analysis of the (2, 1) blocks above, we already know that *Q*_m_ = *Q*_p_.

Thus, using similar arguments, we conclude that 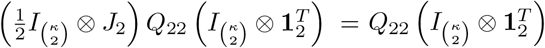.

#### Lumpability criterion

Therefore, for the independent phased process to be lumpable with respect to the unphased partition *G*_u_, the generator matrices of the maternal and paternal processes must be identical. Conversely, if *Q*_m_ = *Q*_p_, then *V*_u_*U*_u_*QV*_u_ = *QV*_u_ is immediate from the symmetry of the generator matrix under interchange of maternal and paternal labels.

The analysis of this section proves the theorem presented below.

##### Theorem 4.2.

*Let X*(*t*) = (*X*_m_(*t*), *X*_p_(*t*)) *be an independent phased maternal-paternal process on the finite state space G with a generator matrix Q*_m_ ⊕ *Q*_p_. *Let G*_u_ *be the unphased partition of the state space as defined by Equation 18. Then X*(*t*) *is lumpable with respect to G*_u_ *if and only if Q*_m_ = *Q*_p_. *In this case, we call the lumped process an* ***independent unphased maternal-paternal process***, *denoted by* 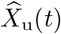.

Using our running example on 9 states (phased diploid ternary) and letting *q*_*i*→*j*_ := *ρ*_*i*→*j*_ = *µ*_*i*→*j*_, the generator matrix for unphased independent process is

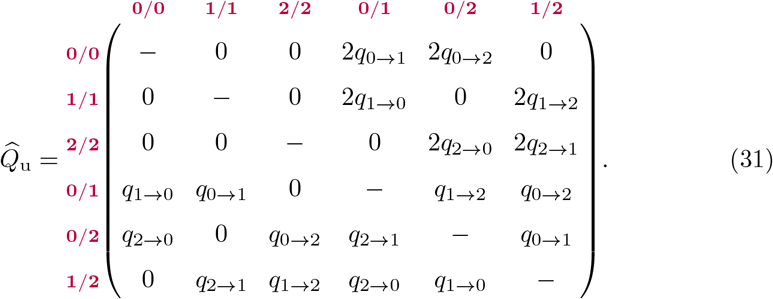

For a graphical representation of this rate matrix, see Figure 1.

#### 4.2.1 A few key points about the independent phased and unphased processes

1. Given a continuous-time Markov process *X*(*t*) = (*X*_m_(*t*), *X*_p_(*t*)) on the state space *G*, where the marginal maternal *X*_m_(*t*) and paternal *X*_p_(*t*) processes are Markovian, the independence of the marginal processes implies that the genotype *i*_m_*j*_p_ cannot simultaneously change both states in an infinitesimal amount of time. Models with this property are referred to as independent phased processes in this article (see Definition 4.1).
2. The unphased partition *G*_u_ as defined by the Equation 18 is a very specific partition that groups together only the states {*i*_m_*j*_p_, *j*_m_*i*_p_}_*i ≠j*_. Of course, other partitions are possible; however, when we refer to an unphased partition, we will always mean the one defined by Equation 18. Lumping of the independent phased model with respect to other partitions is discussed in Section 7.
3. By an independent unphased model, we will always mean the lumped independent phased process with respect to the unphased partition *G*_u_. An independent unphased model necessarily implies that maternal and paternal sequence evolution follow the same Markov process.
4. Our lumpability arguments in this section employed a block form of the generator matrix by permuting rows and columns using a permutation matrix Σ, allowing us to express *V*_u_ (as per Equation 22) in a block form. The relationship between *V*_u_ and its form using lexicographically ordered rows, 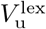, is given by

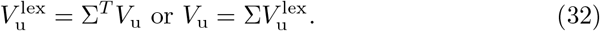

Recall that *V*_u_ and 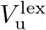 have identical column orderings; the only difference is the ordering of the rows. Similarly, *U*_u_, the left inverse of *V*_u_, maintains its row orderings while its columns are permuted in the 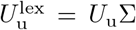 form. Thus, the lumped matrix *Q*_u_ has its rows and columns always ordered in the same manner as the columns of *V*_u_, regardless of whether the blocked form or not was used for the phased generator matrix.

### 4.3 Stationary Distributions

Under the assumption that the independent phased process *X*(*t*) = (*X*_m_(*t*), *X*_p_(*t*)) is irreducible, there is a stationary distribution 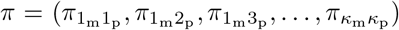, which is also the limiting distribution. Recall that the marginal processes are lumpable. Then, from the discussion of the stationary distribution of the lumped process in Section 3, both the maternal and paternal processes have stationary distributions *π*_m_ = *πV*_m_ and *π*_p_ = *πV*_p_, where *V*_m_ and *V*_p_ are defined as in Section 4.1.1. That is, *π*_m_ and *π*_p_ are obtained by summing the components of *π* in the same partition class:

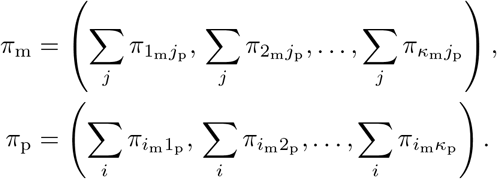

On the other hand, the generator matrix of the independent phased process is a Kronecker sum: *Q*_m_ *Q*_p_. The stationary distribution *π* is a left eigenvector of *Q*_m_ ⊕ *Q*_p_ and its transpose is a right eigenvector of (*Q*_m_ *Q*_p_)^*T*^ both corresponding to the same eigenvalue *λ* = 0.

#### Remark 4.3.

*If λ*_*i*_ *are eigenvalues of matrices A*_*i*_ *for i* = 1, 2, *with corresponding right eigenvectors v*_*i*_, *then λ*_1_ + *λ*_2_ *is an eigenvalue of the Kronecker sum A*_1_ ⊕ *A*_2_ *and v*_1_ ⊗ *v*_2_ *is a corresponding eigenvector, where* ⊗ *denotes the Kronecker product of two vectors. Furthermore, every eigenvalue/eigenvector pair of A*_1_ ⊕ *A*_2_ *arises in this manner. For a proof and additional details see Horn and Johnson [19, Theorem 4*.*4*.*5]*.

This remark implies that each entry 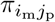 in the stationary distribution *π* is of the form 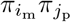, where 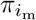 and 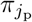 are the corresponding entries of the maternal and paternal stationary distributions *π*_m_ and *π*_p_, respectively, determined by their generator matrices *Q*_m_ and *Q*_p_. Indeed, for all *i* ∈ [*κ*],

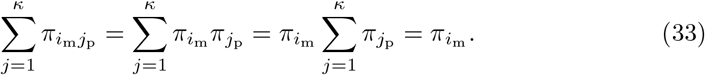

Similarly, the corresponding identity holds for the entry 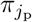.

#### Corollary 4.4.

*Let X*(*t*) = (*X*_m_(*t*), *X*_p_(*t*)) *be an independent phased maternal-paternal process on the finite state space G with a generator matrix Q*_m_ ⊕ *Q*_p_. *Let* 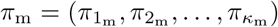 *and* 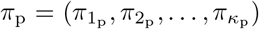 *be stationary distributions for a maternal and paternal processes, respectively. The stationary distribution π of the process X*(*t*) *is given by π* = *π*_m_ ⊗ *π*_p_. *That is*,

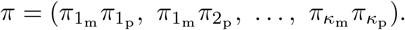

#### Stationary distribution of the independent unphased model

According to Theorem 4.2, an independent unphased maternal-paternal process must have identical generator matrices, *Q*_m_ = *Q*_p_, and thus identical stationary distributions. Let *π*_mp_ = (*π*_1_, …, *π*_*κ*_) denote the stationary distribution of both the maternal and paternal processes. According to Corollary 4.4, the stationary distribution of the independent phased model in this case is *π*_mp_ ⊗ *π*_mp_. The stationary distribution 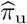 of the independent unphased process is then given by

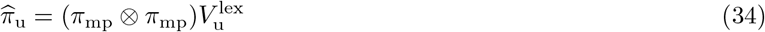

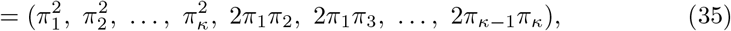

where we use 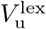 rather than *V*_u_ because the Kronecker product *π*_mp_ *π*_mp_ comes naturally in lexicographic order. For more information, refer to Section 4.2.1, key point (4), and Equation 32. Note that the entries of 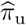 corresponding to classes *i/i* = {*i*_m_*i*_p_} are of the form 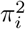, and the entries corresponding to classes *i/j* = {*i*_m_*j*_p_, *j*_m_*i*_p_}_*i*≠*j*_ are of the form 2*π*_*i*_*π*_*j*_.

### 4.4 Time Reversibility of Independent Phased and Unphased Processes

Up to this point, we did not impose the condition that maternal and paternal processes must have time-reversible generator matrices. In this section, we show that the independent phased process and its lumped unphased process are time-reversible if the marginal processes are time-reversible. We begin with a general result.

#### Theorem 4.5.

*Let X*(*t*) *be a continuous-time, time-reversible Markov process on the finite state space with generator matrix Q and unique stationary distribution π. Suppose X*(*t*) *is lumpable with respect to some partition E. Then, the lumped process* 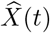 *is time-reversible*.

*Proof* Since the process *X*(*t*) is time-reversible then *D*_*π*_ *Q* = *Q*^*T*^ *D*_*π*_, where *D*_*π*_ = diag(*π*) is a diagonal matrix with *π* on the main diagonal. Moreover, the process is lumpable with respect to partition *E*. Then by Remark 3.4 *V UQV* = *QV* and the generator matrix of the lumped process is 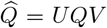, with *V* and *U* as defined in Section 3. The stationary distribution of the lumped process, *πV*, can be represented concisely as the diagonal matrix: 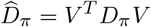.

Then

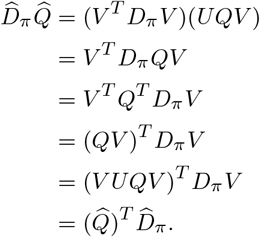

This proves that the lumped process is also time-reversible. □

In light of this theorem, all we need to show is that the generator matrix of the independent phased process is time-reversible, provided that the marginal processes are time-reversible. Thus, assume that *Q*_m_ and *Q*_p_ are time-reversible, meaning that

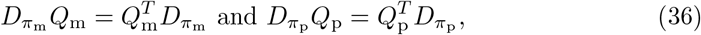

where 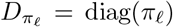 for ℓ m, p. It is immediate that an independent phased process is also time-reversible. To see this, note that

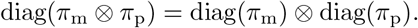

Therefore,

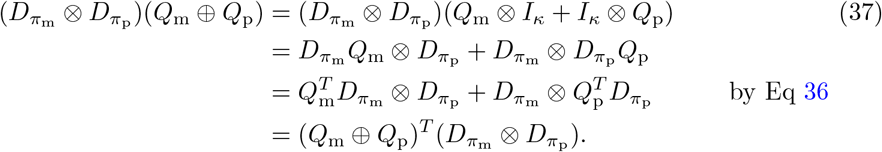

The above discussion, combined with Theorem 4.5, proves the following result.

#### Lemma 4.6.

*If marginal maternal and paternal processes are time-reversible, then both the independent phased process and its lumped unphased process are also time-reversible*.

### 4.5 Symmetric matrix of the time-reversible phased and unphased processes

Under the assumptions of the models discussed here (e.g., existence of the limiting distribution) and the time-reversibility of the process, the generator matrix can be expressed as a product of two symmetric matrices. In particular, if *Q*_m_ and *Q*_p_ are time-reversible, then 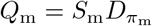 and 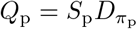, where *S*_m_ and *S*_p_ are symmetric matrices and 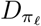 is defined as above, ℓ ∈ {m, p}. This decomposition leads to the traditional expression *π*_*j*_*r*_*i↔j*_, *i* ≠ *j*, of the off-diagonal entries of the generator matrix. Using our running example on 9 states, a time-reversible maternal generator matrix *Q*_m_ is

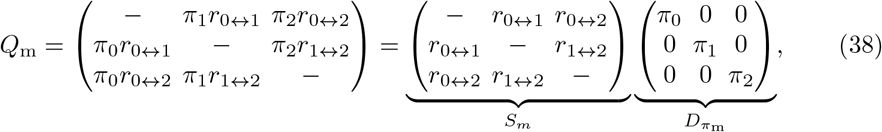

with diagonal entries such that the rows of *Q*_m_ sum to zero. The parameters *r*_*i↔j*_ are called exchangeability parameters and the symmetric matrix *S*_*m*_ is the exchangeability matrix. The paternal case has an identical form. The symmetric part for the independent phased model can be computed as follows:

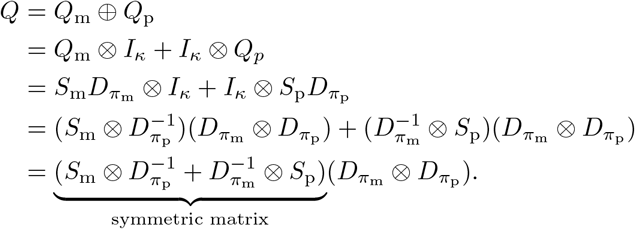

This demonstrates that the symmetric part of the generator matrix for an independent phased process can be expressed in terms of the stationary distributions and symmetric matrices for the maternal and paternal processes. We denote this matrix as

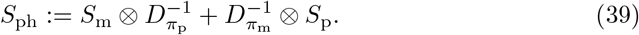

To show that *S*_ph_ is symmetric is a straightforward exercise.

Now we compute the symmetric part of the independent unphased process. Since *Q*_m_ = *Q*_p_, let *Q*_mp_ denote the common generator matrix for both processes. To find the symmetric matrix of the unphased rate matrix we will need lemma below, which is proved in Appendix A.

#### Lemma 4.7.

*Let X*(*t*) *be a continuous-time Markov process on a finite state space S with generator matrix Q and unique stationary distribution π. Suppose X*(*t*) *is lumpable with respect to a partition E of S with r classes. Then*,

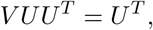

*where V and U are as defined in Section 3. Moreover, if π is constant on each partition class, then*

1. *D*_*π*_*V U* = *V UD*_*π*_, *where D*_*π*_ = diag(*π*), *and*
2. 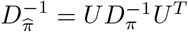, *where* 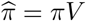 *and* 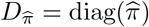.

Recall that the stationary distribution 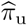 of the independent unphased process is given by

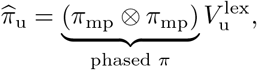

where *π*_mp_ is the stationary distribution for both the maternal and paternal processes. To simplify the notational burden in computing the symmetric part of the unphased rate matrix, we will omit the subscripts and superscripts from 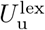 and 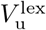, and the subscript from 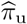. For the computations below, we do not need the expanded representation, but for completeness keep in mind that *Q* = *Q*_mp_ ⊕ *Q*_mp_ and *D*_*π*_ = diag(*π*_mp_ ⊗ *π*_mp_).

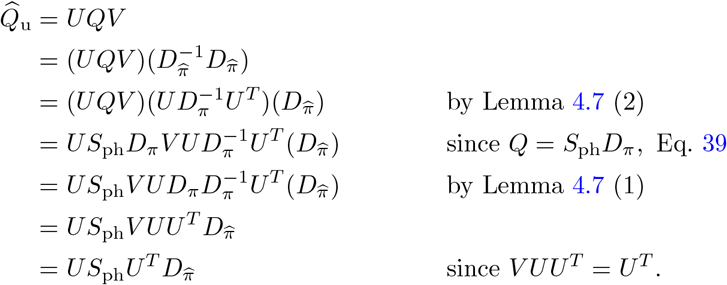

Therefore, the unphased generator matrix 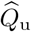 can be written in the form

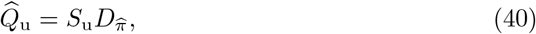

for the symmetric matrix *S*_u_ = *US*_ph_*U*^*T*^.

### 4.6 Scaling of the Independent Phased and Unphased Generator Matrices

As mutation rates can vary over time, it is common to measure the branch lengths of phylogenetic trees in terms of the expected number of mutations rather than calendar (clock) time. Furthermore, for continuous-time Markov chains, there is a relationship between an overall scaling factor of the generator matrix and branch lengths. That is, given a scaling factor *µ >* 0, the Markov matrices *P* (*t*) = exp(*Qt*) and *P* (*t*_∗_) = exp(*Q*_∗_*t*_∗_) are equal when 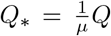 and *t*_∗_ = *µt*. As such, generator matrices are usually assigned a fixed normalization to resolve this non-identifiability. To derive the general scaling factor for the models discussed in this article, we first make a few convenient definitions.

#### Definition 4.8.

*Let Q be a generator matrix and π a stationary distribution (as a row vector). Let d be the row vector whose entries are diagonal entries of Q. The scaling factor µ is defined as*

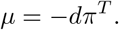

*The normalized form of Q is then simply* 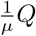.

Now, the independent phased model and each of its maternal and paternal submodels have their own *µ*, and we must ask what relationships exist among these quantities. Without this information, it is not possible to appropriately scale inferred branch lengths in a phylogenetic tree when, for example, sequences are simulated under independent maternal and paternal processes, but inference is performed at the level of the phased (or unphased) process.

Recall, that the generator matrix for the independent phased model is *Q* = *Q*_m_ ⊗ *I*_*κ*_ + *I*_*κ*_ ⊗ *Q*_p_. The diagonal entries of each summand, when viewed as row vectors, are then

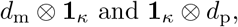

where **1**_*κ*_ is a row vector of all 1’s of length *κ*. The scaling factor for the independent phased process is now easily computed:

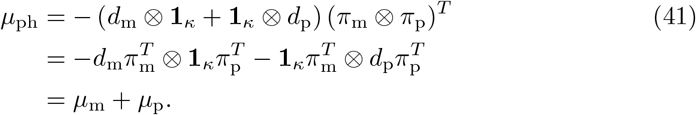

The conclusion in the above equation follows from the fact that 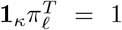 for ℓ ∈ {m, p}. If *Q*_m_ = *Q*_p_ with scaling factor *µ*_mp_, then *µ*_ph_ = 2*µ*_mp_.

From the perspective of branch length inference, this implies that we should optimize the generator matrix for the independent phased model subject to the constraint *µ*_ph_ = 2 when the maternal and paternal sequences are generated using rate matrices normalized so that *µ*_m_ = *µ*_p_ = 1.

In general, we do not expect the scaling factor of a lumped process to be the same as that of the corresponding unlumped process. However, like the independent phased process, the scaling factor of the independent unphased process is also 2*µ*_mp_. Intuitively, this is because *µ* records the expected number of mutations per unit time and, if there are non-zero mutation rates between states within the same lumped class, these contribute to the rate of the unlumped model but not to that of the lumped model. Following this intuition, since the transition rates between mixed heterozygous states *i*_m_*j*_p_ → *j*_m_*i*_p_ are all zero in the phased model, there is no difference between the *µ* values for the phased and unphased models.

More concretely, let *Q*_mp_ := *Q*_m_ = *Q*_p_, and let *d*_mp_ denote the row vector of diagonal entries of *Q*_mp_. As before, we drop the subscripts and superscripts from 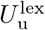 and 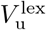, but keep in mind that what follows requires the states to be ordered lexicographically.

First note that

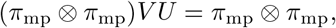

since *V U* averages within partition classes and *π*_mp_ *π*_mp_ is constant on each class.

Additionally, the diagonal entries of the two summands in

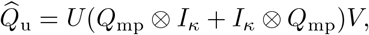

viewed as row vectors, are

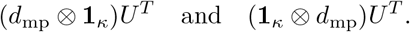

Then the scaling factor *µ*_u_ of the independent unphased process is

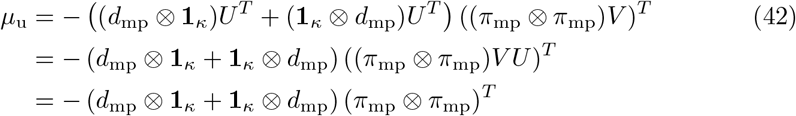

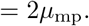

As before, from the perspective of branch length inference, this implies that we should optimize the generator matrix for the independent unphased model subject to the constraint *µ*_u_ = 2 when the maternal and paternal sequences are generated using rate matrices normalized so that *µ*_m_ = *µ*_p_ = 1.

## 5 Likelihood Factorization

Likelihood methods are central to phylogenetic inference, and to present our ideas clearly, we focus on the computation over a specific tree and state space under an independent genotype model. We make two common assumptions: sites evolve independently, and lineages evolve independently. Additionally, we use Felsenstein’s pruning algorithm [20] to demonstrate the computation of site pattern probabilities, but with a slight difference in notation.

Given an internal node *s* of a rooted tree, the usual formulation of Felsenstein’s pruning algorithm [20] writes the probability distribution of site patterns at *s* in terms of a product of the likelihoods of the subtrees hanging off of *s*, conditional on the state at *s*. While this approach gives a straightforward formula in components, for our purposes it is more convenient to express the same computation using the pointwise (Hadamard) product of matrices, denoted by ⊙. This notation has the helpful property that, if *A*_1_ and *A*_2_ are matrices of the same dimension, and *B*_1_ and *B*_2_ are also matrices of the same dimension, then the mixed Hadamard-Kronecker product satisfies

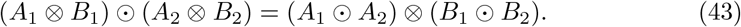

Another mixed-product property that we will utilize relates the Kronecker product and matrix multiplication:

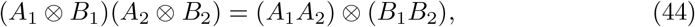

where *A*_*i*_ and *B*_*i*_ are such that the products *A*_1_*A*_2_ and *B*_1_*B*_2_ are defined.

These relations are useful because in our setting the transition probability matrix associated with each branch of the tree is a Kronecker product of two transition matrices corresponding to maternal and paternal Markov processes.

### Site-pattern probability

Consider a rooted three-taxon tree *T* = ((1:*t*_1_, 2:*t*_2_):*t*_4_, 3:*t*_3_), as depicted in Figure 2A. Let *X*_ℓ_ denote the state observed for lineage ℓ = 1, 2, 3, and let *P* (*t*_*i*_) denote the transition probability matrix associated with edge *t*_*i*_. The joint probability

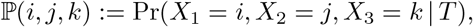

also referred to as the site pattern probability, is given by

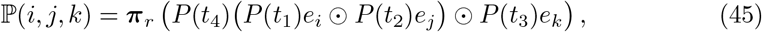

where ***π***_*r*_ is the probability distribution of states at the root and *e*_*s*_ is the standard basis vector in ℝ^*n*^ with a 1 in the position corresponding to the state *s* ∈ {1, 2, …, *n*}. Details are provided in Appendix B. Even though the taxa in Figure 2A are labeled by genotypes, the expression in (45) holds for an arbitrary finite-state Markov model.

**Fig. 2:**
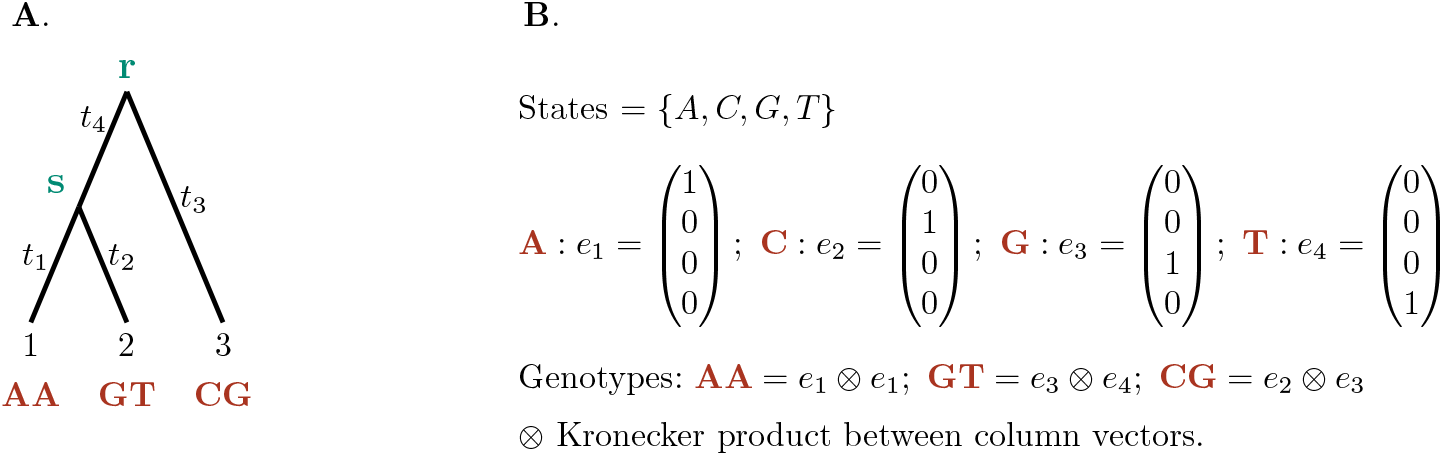
(**A**) A three-taxon binary tree with arbitrary branch lengths *t*_*i*_ ≥ 0. The genotype data at a single site is depicted below the leaves. (**B**) Assignment of standard vectors (probability vectors) to nucleotide bases. Genotypes are the Kronecker product between the two standard vectors.

### Likelihood for independent phased genotype model

Using Equation 45, we first demonstrate the computation of a site pattern probability under the phased genotype state space

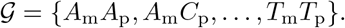

This example illustrates how site pattern probabilities factor into maternal and paternal components. The same factorization naturally extends to arbitrary trees under the independent phased genotype model, leading to a corresponding factorization of the likelihood.

To simplify the notation, the maternal and paternal subscripts (*m* and *p*) will be omitted from genotypes; for example, *G*_m_*T*_p_ will be written simply as *GT*. The ordering of alleles is unchanged. Assume independent phased maternal and paternal Markov processes across the branches of the tree in Figure 2A, with root distribution given by the row vector ***π***_*r*_ = *π*_m_ ⊗ *π*_p_ and transition probability matrix *P*_m_(*t*_*k*_) ⊗ *P*_p_(*t*_*k*_) on each branch *t*_*k*_. To simplify the notation further, let *M*_*k*_ := *P*_m_(*t*_*k*_) and *F*_*k*_ := *P*_p_(*t*_*k*_). Thus, *M*_*k*_ and *F*_*k*_ denote the *maternal* and *paternal* transition probability matrices, respectively, associated with branch *t*_*k*_. Associate a standard column vector *e*_*i*_ ∈ ℝ^4^ to each nucleotide, allowing phased genotypes to be represented as Kronecker products of the form *e*_*i*_ ⊗ *e*_*j*_, where *i, j* ∈ {1, 2, 3, 4} (Figure 2B). Note that

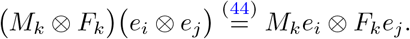

Using Equations 43 and 44 repeatedly, the site pattern probability ℙ (*σ*), where *σ* = (*AA, GT, CG*), is computed by direct substitution into Equation 45:

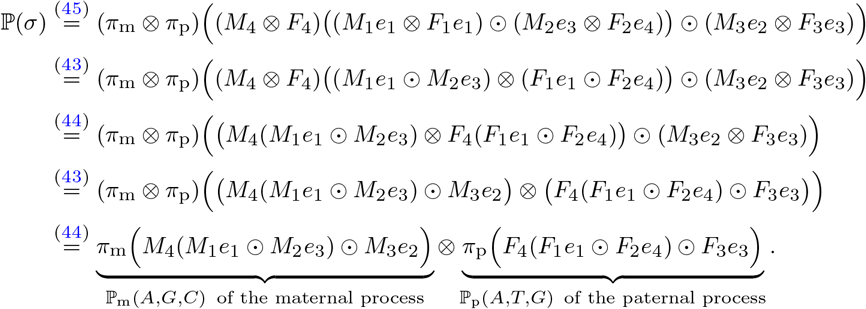

Since the Kronecker product of 1 *×* 1 matrices is simply the ordinary scalar product, the joint probability factors as

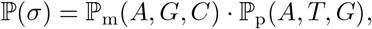

where ℙ_m_(*A, G, C*) and P_p_(*A, T, G*) are the site pattern probabilities of the maternal and paternal processes, respectively.

In general, let *σ* = (*g*_1_, *g*_2_, *· · ·, g*_*n*_) denote a phased genotype site pattern on a fixed tree relating *n* cells, where *g*_*k*_ = *i*_*k*_*j*_*k*_ is the phased genotype observed at taxon *k*, with maternal allele *i*_*k*_ and paternal allele *j*_*k*_. Then the probability of genotype pattern *σ* under the proposed independent phased genotype model on the fixed tree satisfies

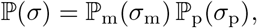

where *σ*_m_ = (*i*_1_, …, *i*_*n*_) and *σ*_p_ = (*j*_1_, …, *j*_*n*_) are the corresponding maternal and paternal allele site patterns on the same tree.

Then, under the assumption of site independence, the likelihood is

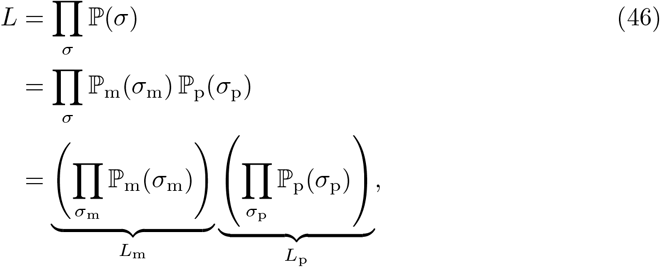

where we let *σ* range over all the sites in the alignment. Here *L*_m_ and *L*_p_ denote the maternal and paternal likelihoods, respectively. Of course, in practice identical site patterns are grouped and weighted by their observed frequencies.

In general, even if *σ*_m_ = *σ*_p_, ℙ_m_(*σ*_m_) ≠ ℙ_p_(*σ*_p_), unless the maternal and paternal Markov processes are identical on the tree. This means one has to be careful when aggregating maternal and paternal patterns. For example, consider the genotype patterns

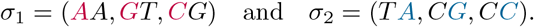

When the maternal and paternal processes are different, the site pattern probabilities

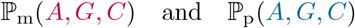

contribute differently and therefore should not be combined.

### Likelihood for independent unphased genotype model

Using our running toy example in Figure 2, we first show that the site pattern probability under the independent unphased model defined in Section 4.2 is a sum of phased site pattern probabilities. Under the unphased model, the phased site pattern (*AA, GT, CG*) in Figure 2A is represented by *σ*_u_ = (*A/A, G/T, C/G*). Recall that the independent unphased genotype model necessarily has identical maternal and paternal processes (Theorem 4.2), implying that *M*_*k*_ = *F*_*k*_ and *π*_m_ = *π*_p_. Let the common transition probability matrix and root distribution be denoted by *M*_*k*_ and *π*, respectively. In particular, let

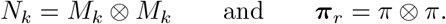

Associated with each branch *t*_*k*_ is the lumped transition probability matrix

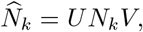

where the rows of *U* and columns of *V* are ordered lexicographically (see the discussion in Section 4.2.1). For cleaner notation below, we omit both subscripts and superscripts from these matrices whenever no ambiguity arises. The root distribution is the row vector ***π***_*r*_*V*. Finally, let *e*_*ij*_ = *e*_*i*_ ⊗ *e*_*j*_. Then unphased genotypes are represented by the column vectors *V* ^*T*^ *e*_*ij*_.

The computations below rely on the following identities, whose proofs are given in Appendix A:

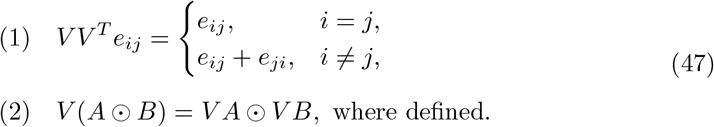

The first identity is specific to the *V* matrix for the unphased partition. The second identity also does not hold for an arbitrary matrix *V* and is a special property of *V* matrices defined in the construction of lumped processes.

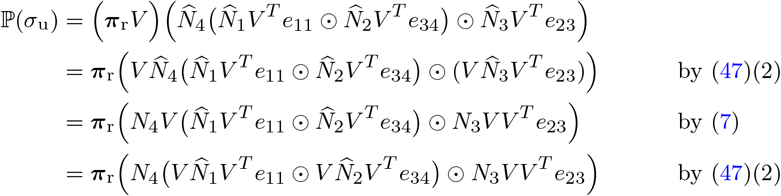

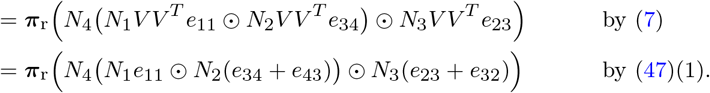

Distributing Hadamard product and root distribution over the sums results in

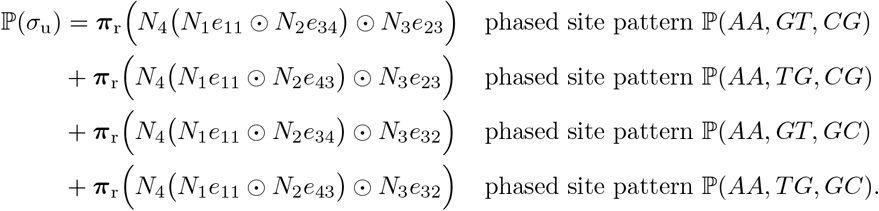

It is clear that each line in the equation above is a phased site pattern probability corresponding to a phased pattern consistent with *σ*_u_ = (*A/A, G/T, C/G*) with each heterozygous genotype contributing two possible maternal-paternal assignments. Recalling that a phased site pattern probability factors into maternal and paternal site pattern probabilities, we obtain

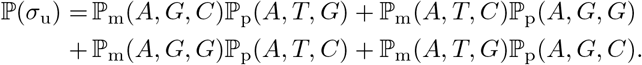

Note the distinction between phased and unphased site pattern probabilities. Although the unphased site pattern probability is obtained by summing the corresponding phased site pattern probabilities, the maternal and paternal processes are identical under the independent unphased model. This means that, ℙ_m_(*σ*_m_) = ℙ_p_(*σ*_p_) whenever *σ*_m_ = *σ*_p_. To simplify the notation, we therefore omit the subscripts m and p from site pattern probabilities in the remainder of this section and write ℙ_m_(*σ*_m_) = ℙ (*σ*_m_) and ℙ_p_(*σ*_p_) = ℙ (*σ*_p_).

In general, let [*σ*_u_] denote the equivalence class of phased genotype patterns consistent with the unphased pattern *σ*_u_. Then the probability of unphased genotype pattern *σ*_u_ under the proposed independent unphased model on a fixed tree is

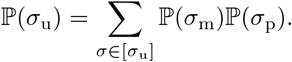

Consequently, the likelihood under the independent unphased model is

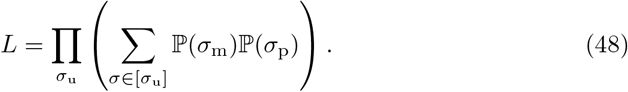

Mathematically, the expression above establishes a direct connection between the likelihoods of the phased and unphased models. Computationally, however, this representation is generally impractical for large single-cell datasets. Since each heterozygous genotype contributes two possible maternal-paternal assignments, an unphased site pattern with *h* heterozygous taxa corresponds to 2^*h*^ phased site patterns. Now if alignment contains hundreds of cells and thousands of unphased genotypes, the number of underlying phased site patterns can become prohibitively large.

## 6 Some Dependent Models

In Sections 4.1 and 4.2, we considered diploid models based on an independence assumption for the haploid maternal and paternal processes. From a mathematical perspective, it is then natural to ask how diploid evolution behaves when the maternal and paternal components are dependent. A full treatment of dependence is beyond the scope of the current article and would include classifying models that may have limited biological relevance. Instead, we give two examples that exhibit some of the behaviors arising from dependence and that may have biological interpretations, along with one example that retains desirable mathematical properties despite the presence of dependence.

### 6.1 Dependent non-Markovian Marginal Processes

#### Allelic interaction model

Consider a model in which the maternal and paternal mutation rates are equal but where the presence of a mutant allele on one of the maternal-paternal strands affects the mutation rate on the other strand. For example, on the phased *κ* = 2 state space, a possible rate matrix for such a process is

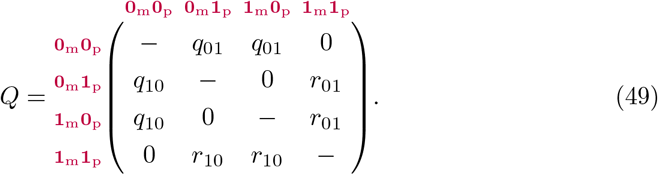

where *q*_01_ ≠ *r*_01_ and *q*_10_ ≠ *r*_10_.

One can check that this model is not lumpable with respect to the maternal-paternal partitions. In particular, since we assume both (i) non-independence of the maternal and paternal components and (ii) that the rates of simultaneous changes are zero, the marginal processes cannot be Markovian. The latter is an implication of Ball and Yeo [18, Theorem 3.2], discussed in Section 4.1.

However, this model is still unphased-lumpable with respect to the unphased partition defined in Equation 18: *G*_u_ = {0/0, 1/1, 0/1}. The unphased lumping gives the matrices

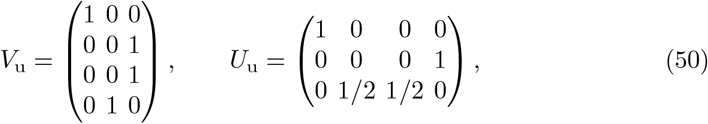

and it is easy to check that *V*_u_*U*_u_*QV*_u_ = *QV*_u_. Hence, the model is unphased-lumpable and the lumped model is

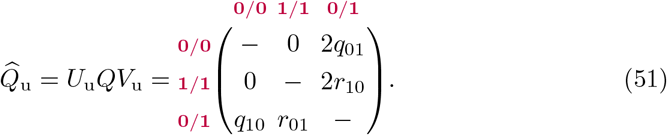

#### Mitotic recombination model

To form a basic model that incorporates mitotic recombination, we can begin with the phased independent maternal-paternal model and add a rate for transitions between phases. It is important to note that this addition breaks the independence of the maternal and paternal processes since, for example, point mutations in the maternal DNA depend only on the current maternal state but, if a recombination event occurs, the result also depends on the paternal state. However, such a model is still unphased-lumpable and, in fact, has the same unphased lumping as the recombination-free version. For example, consider the *κ* = 2 independent phased model with generator matrix *Q* = *Q*_mp_ ⊕ *Q*_mp_ satisfying unphased lumpability constraint, i.e., *Q*_mp_ = (*q*_*ij*_). Let *r >* 0 be the phase transition rate between genotypes 0_m_1_p_ and 1_m_0_p_. The rate matrix for the corresponding recombination model is

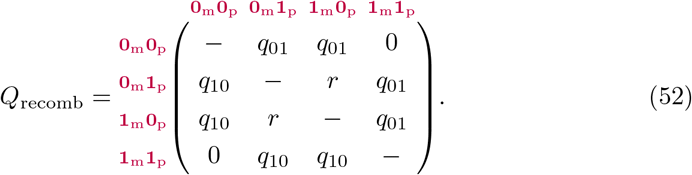

Similar to our first example, this model is not lumpable with respect to the maternal-paternal partitions, but it is lumpable with respect to unphased partition.

This particular model can be defined compactly for any number of states. First, note that the generator matrix given by Equation 52 is the sum of two matrices:

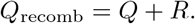

Second, using the block-adapted ordering of the phased state space, *R* can be written in block-diagonal form:

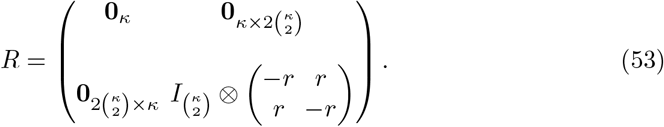

To show that this model is consistent with an unphased lumping, we must verify the lumping condition *V*_u_*U*_u_(*Q* + *R*)*V*_u_ = (*Q* + *R*)*V*_u_, with *V*_u_ and *U*_u_ as in Equation (24).

Now, we have already established that *V*_*u*_*U*_*u*_*QV*_*u*_ = *QV*_*u*_ in Section 4.2, so we only need to check that *V*_*u*_*U*_*u*_*RV*_*u*_ = *RV*_*u*_. The block structure of *V*_*u*_ and *R* is such that the only non-trivial matrix multiplication is

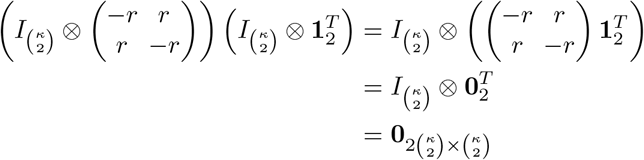

This means that *RV*_u_ = **0** and so *V*_u_*U*_u_*RV*_u_ = *RV*_u_ is satisfied. In fact, we have shown something stronger. Not only is *Q*_recomb_ lumpable with respect to the unphased partition, but the unphased lumped models for *Q*_recomb_ and *Q* = *Q*_mp_ ⊕ *Q*_mp_ are identical, since *U*_u_*Q*_recomb_*V*_u_ = *U*_u_(*Q* + *R*)*V*_u_ = *U*_u_*QV*_u_ = *Q*_u_.

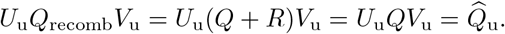

### 6.2 Dependent Markovian Marginal Process

The previous two examples illustrate dependence arising from interaction between maternal and paternal evolution and from recombination-like phase-switching events. A model with Markovian marginals can be obtained from the mitotic recombination model; however, the biological interpretation of this model is less clear. Nevertheless, it provides an example of a dependent process with Markovian maternal and paternal marginals.

The model below is lumpable with respect to all three partitions: the maternal, paternal, and unphased partitions.

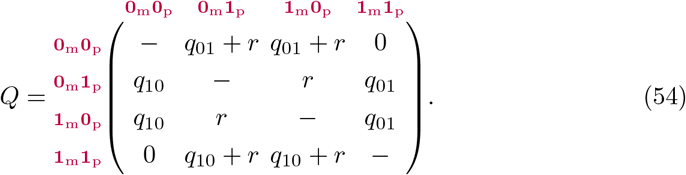

With respect to the unphased partition the lumped generator matrix is:

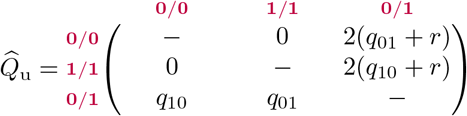

Unlike the mitotic recombination model, this lumped model is not the same as the unphased independent model. Additionally, its marginal processes, which are identical for the maternal and paternal partitions, are of the form *Q*_mp_ plus a recombination-like term:

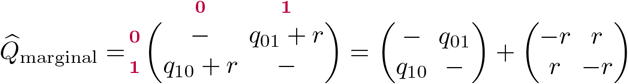

The construction of this model is not ad hoc and extends to an arbitrary number of states. The resulting family has several interesting mathematical properties. A detailed study of these models is beyond the scope of the present article and will be pursued in subsequent work.

## 7 Some Models from the Literature

The models selected for this section are all finite-sites models of genotype evolution defined on either phased or unphased genotype spaces. While these models differ in some of their assumptions, two of them assume that a genotype cannot change both of its states simultaneously in an infinitesimal amount of time. The latter assumption is one of the three conditions in our definition of an independent phased process (see Definition 4.1), but it is not sufficient for independence. One model, CellPhy [12], makes the additional assumption that maternal and paternal chromosomes evolve independently and that transition rates are independent of the homologous allele; nevertheless, the model as presented cannot be obtained from a truly independent phased process. The same is true for the other models.

### SiFit

Models presented in Zafar et al. [8] are defined directly over three states or two states. The three-state model is specific to diploid data that differentiates between homozygous and heterozygous mutations. They define their states as: 0 = homozygous reference, 1 = heterozygous, and 2 = homozygous non-reference. So that the states of the (ternary) SiFit3 model correspond to the following unphased states:

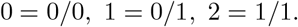

The SiFit3 model, as defined in Zafar et al. [8, Equation (5)], cannot be the result of the lumped independent phased model with respect to the unphased partition, as can be seen in the matrices below:

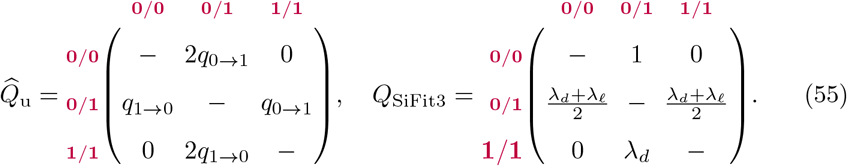

The parameters *λ*_*d*_ and *λ*_ℓ_ of the generator matrix *Q*_SiFit3_ represent deletion and loss of heterozygosity, respectively. One can see that the equalities

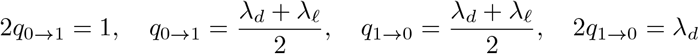

cannot be simultaneously true unless *λ*_*d*_ = 1 and *λ*_ℓ_ = 0, which then implies that *q*_0→1_ = *q*_1→0_ = 1/2. On the other hand, *Q*_SiFit3_ can be obtained from the phased dependent model below:

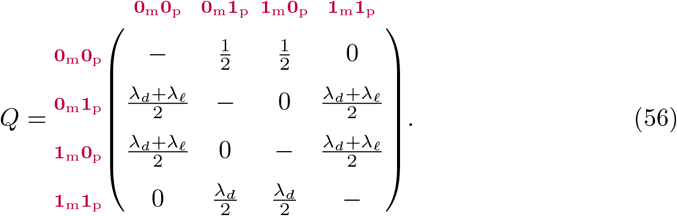

This model is lumpable with respect to the unphased partition, and the lumped generator matrix is *Q*_SiFit3_. The model is not lumpable with respect to the maternal or paternal partition, and hence the marginal processes are not Markovian. The model presented in Equation 56 is one of the general forms and is not unique. The rates between the states 0_m_1_p_ ↔ 1_m_0_p_ can take any value and do not contribute to the rates of the lumped generator matrix. The other constraints are

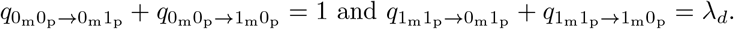

We chose the rates for the phased matrix (56) above to match closely the spirit of the SiFit model.

The state space of the binary model in SiFit is defined as absence (0) or presence (1) of a mutation. While no further information is given regarding how the binary states are produced from the underlying genotypes, the article clearly states that homozygous non-reference and heterozygous genotypes represent mutations. Thus, if the genome is diploid, one can think of collapsing genotypes 1 and 2 into a single class or, more generally, collapsing 01, 10, and 11 into a single class representing the mutation state. In this setting, one can show, similarly to the discussion of the ternary model, that the binary generator matrix is neither the result of a lumped independent phased model nor the result of lumping the ternary SiFit model. A general matrix does exist that is lumpable with respect to the partition {{00}, {01, 10, 11}}, has non-Markovian marginals, and yields the binary SiFit model, but it differs from the matrix in Equation 56. We omit the details here for the following reason. Zafar et al. [8] analyze a binary metastatic colorectal cancer dataset that contains aneuploid cells. In this case, interpreting the state space in terms of diploid genotypes and passing to a phased model becomes unclear. Therefore, we refrain from making any claims about the binary SiFit model.

#### SIEVE

The method behind SIEVE [11] jointly infers somatic variants and cell phylogeny. The generator matrix is defined directly over four classes, {0/0, 0/1, 1/1, 1/1^*′*^}, where 0 represents the reference allele and 1 and 1^*′*^ represent the first and second alternate alleles, respectively. The authors refer to 0/0 as wildtype, 0/1 as a single mutant, 1/1 as a homozygous mutant, and 1/1^*′*^ as a double heterozygous mutant. Each class is an aggregation of unphased genotypes, e.g., 0/1 = {0/1, 0/1^*′*^}, which can also be viewed as partitions of the phased state space on nine states:

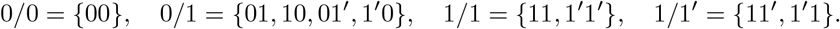

The authors of SIEVE state that at most one change is allowed to occur between their classes; that is, the rate from 0/0 to 1/1 is set to 0. This means that a transition from class 0/1 to class 1/1 is allowed, as depicted in their rate matrix. The claim, however, does not extend to representatives of these classes. For example, the double transition from 01^*′*^ ∈ {01, 10, 01^*′*^, 1^*′*^0} to 11 ∈ {11, 1^*′*^1^*′*^} is potentially allowed to be nonzero. This matters because a lumped generator matrix sums the rates of the corresponding unlumped generator matrix over states belonging to the same class.

Also, a single change between classes does not imply independence of maternal and paternal evolution. To determine whether the model in SIEVE could have originated from an independent phased process, we first derive the conditions under which the matrix given by Equation 10 is lumpable with respect to the SIEVE partition

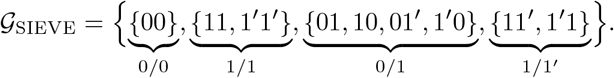

Solving the linear equations arising from the lumpability condition *V UQV* = *QV*, we find that the solutions are restricted to the form

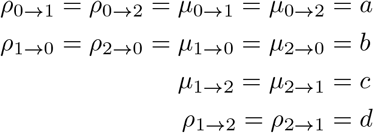

for some constants *a, b, c*, and *d*. Note that the maternal and paternal processes need not be equal since *c* and *d* could differ. The lumped rate matrix *Q* of the independent phased model under these restrictions and the rate matrix *Q*_SIEVE_ from SIEVE are displayed below:

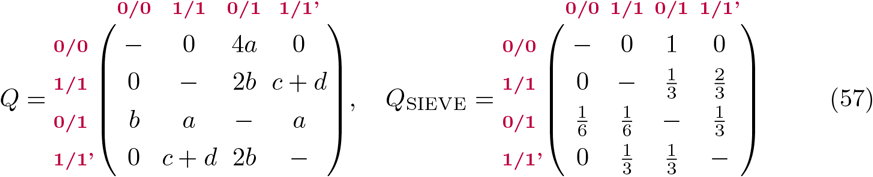

It is easy to see that these two matrices cannot be reconciled for any values of *a, b, c*, and *d*. For example, *Q*_32_ = *Q*_34_ = *a*, but the corresponding entries in *Q*_SIEVE_ are 1/6 and 1/3, respectively. Therefore, the SIEVE model is not the result of lumping an independent maternal-paternal process.

One can obtain a dependent phased model (Equation 58) that is lumpable with respect to the partition *G*_SIEVE_. We omit the details, but the model below is not unique and comes from a general family of lumpable models under this partition.

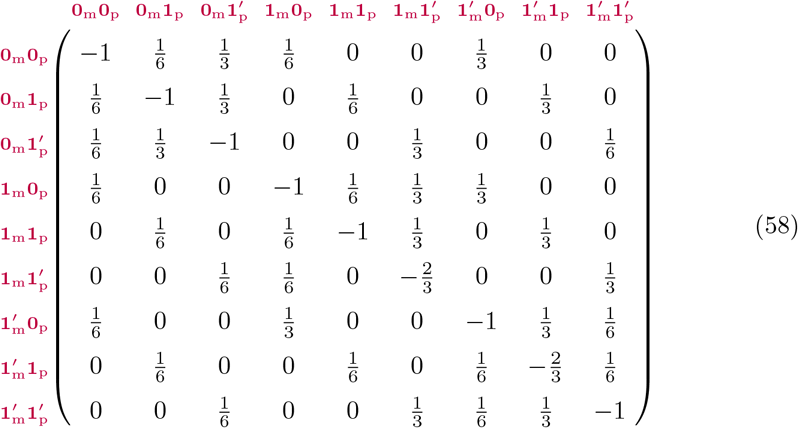

This family contains models that both allow and prohibit double changes in a single time unit. The model presented here is one that restricts phased genotypes from changing both of their states simultaneously. The marginal processes of this model are not Markovian.

#### Cellphy

The GT16 and GT10 models, introduced in Kozlov et al. [12], are time-reversible models based on the phased and unphased DNA state spaces, respectively. The GT16 model is highly similar to the independent maternal-paternal phased model developed in Section 4.1. However, there are notable differences in the stationary distributions. In the GT16 model, the stationary distribution *π*_16_ is expressed in terms of the various *π*_*ij*_ where *π*_*ij*_ need not be the same as *π*_*ji*_ when *i* ≠ *j*. In the independent maternal-paternal phased model, *π*_*ij*_ = *π*_*i*_*π*_*j*_ as in Corollary 4.4. If this relation is added to the GT16 model, it becomes identical to the independent maternal-paternal model. A similar difference occurs between the GT10 model and the unphased independent lumped model. That is, in the unphased model, the stationary distribution is a result of lumping the phased stationary distribution. In GT10, the stationary distribution *π*_10_ has no such constraint.

As discussed in Section 4.4, time-reversible model can be expressed as the product of a symmetric matrix and a diagonal matrix of state frequencies. For the GT10 model, this means that 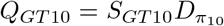 where

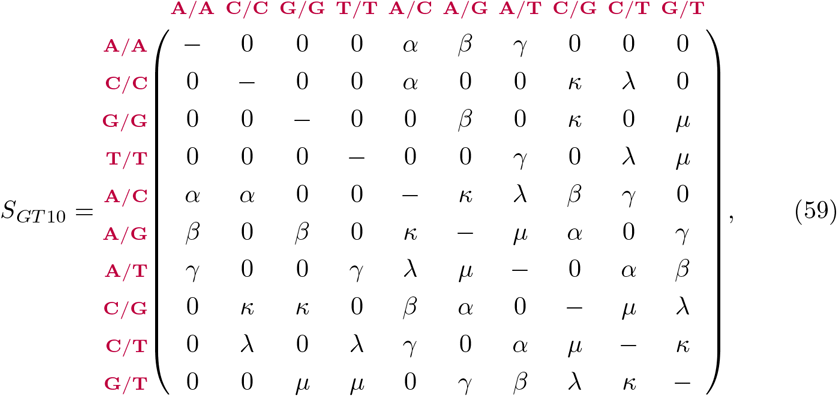

*α, β, γ, κ, λ, µ* are non-negative parameters, and 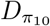 is the diagonal matrix of unphased site pattern frequencies, estimated directly from 10 state counts. In contrast, consider the unphased independent lumped DNA model 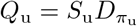, where the underlying maternal and paternal processes are of the form 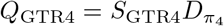 with

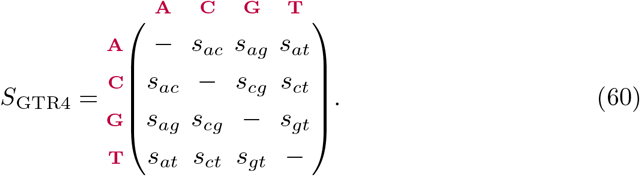

Then, using Equations (39) and (40), we can derive a formula for *S*_u_:

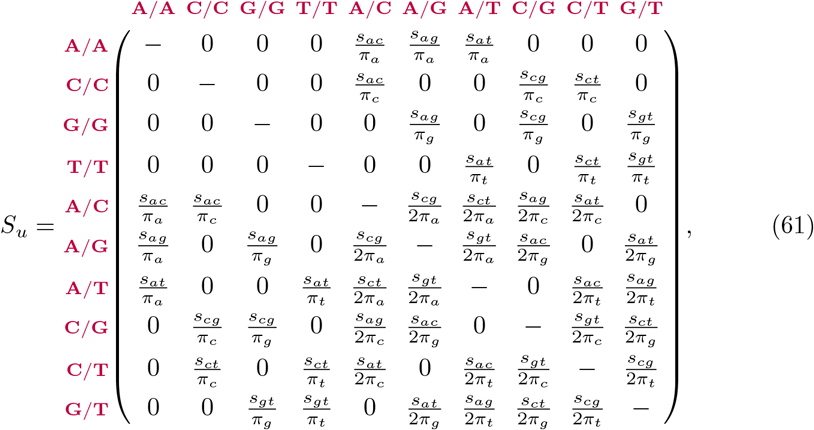

where *π*_4_ = (*π*_*a*_, *π*_*c*_, *π*_*g*_, *π*_*t*_) is the stationary vector for the 4 state DNA model. There are several similarities between the matrices of (59) and (61): both have zeros at double transitions and both have six rate parameters. However, the unphased lumped model both contains DNA frequencies in *S*_u_ and, perhaps more importantly, there are factors of 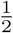 in *S*_u_ (and 2 in *Q*_u_) which appear in a block structure. We will return to this structure in Section 8, where the effect of these factors can be seen in Figure 3. Also note that there are 10 frequency parameters in the GT10 model, one for each of the possible unphased states, while there are only 4 frequency parameters in the unphased independent lumped model; the unphased state frequencies are derived from the base frequencies.

**Fig. 3:**
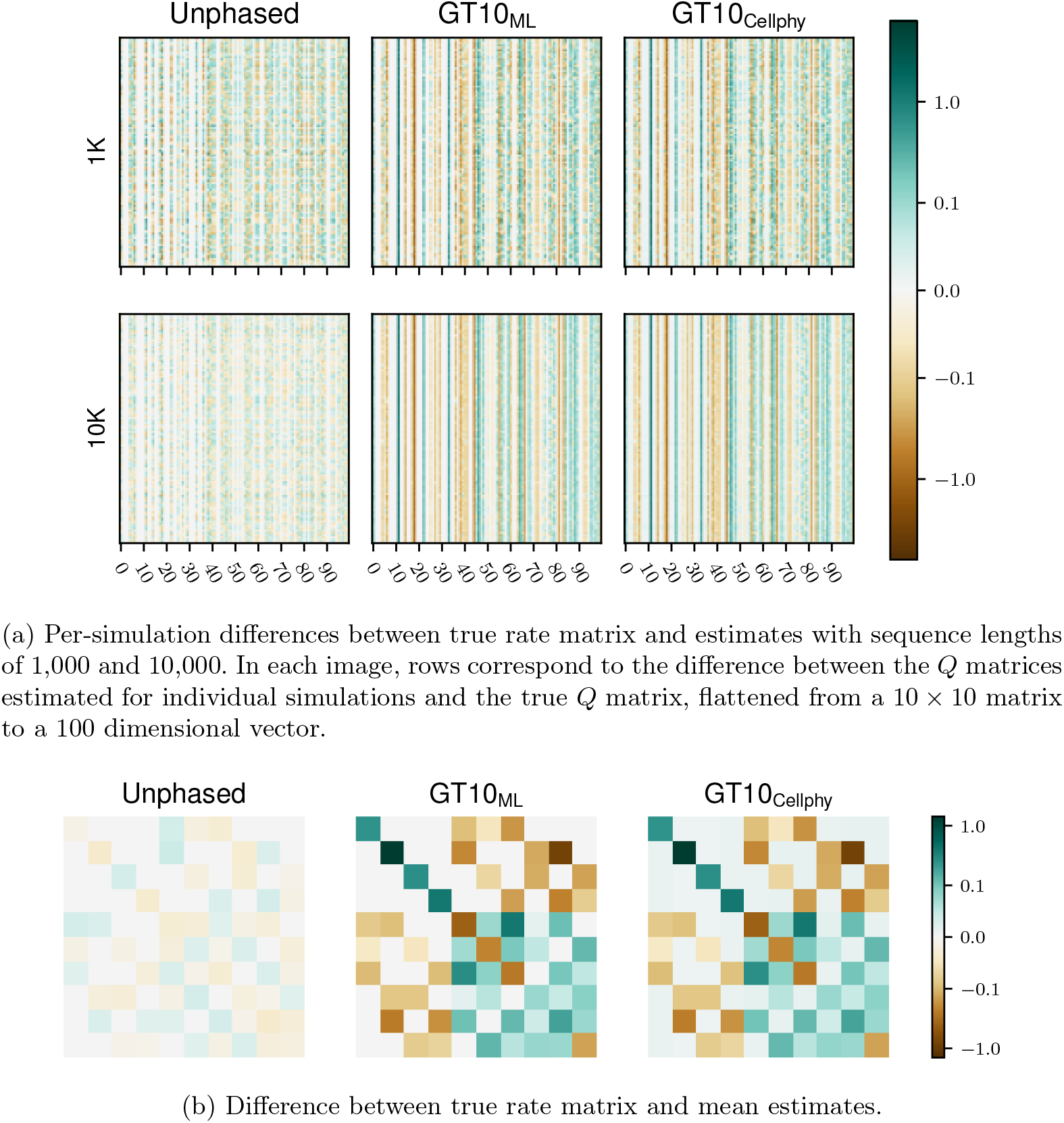
Differences between the estimated rate matrices and the true rate matrix for the 10 state models, with estimates made under the unphased lumped model, the GT10 model estimated using our code (GT10_ML_), and the GT10 model estimated by RAXML-NG (GT10_Cellphy_). Note that both estimated GT10 models are similar, systematically underestimating the rates of homo/heterozygous transitions and overestimating the rates of hetero/heterozygous transitions. Also note that RAXML-NG does not set the rates of double transitions to zero, but instead fixes these rates at 0.001, which accounts for a slight rate overestimate in some entries of the GT10_Cellphy_ estimate.

In fact, although we omit the proof here, it is possible to show something stronger. There is no phased DNA model that is both unphased lumpable and time-reversible which results in the GT10 model. However, there are phased DNA models which are unphased lumpable but which are *n*ot time-reversible that lump to the GT10 model. Importantly, the stationary distribution for the phased 16-state model encodes the stationary distribution for the GT10 model. The general form for the phased 16-state model rate matrix, compatible with GT10, is given in Appendix C. Except when the GT10 model is degenerate, none of these 16 state models are lumpable with respect to marginal maternal and paternal processes implying that the 16-state model is maternal-paternal dependent.

## 8 Simulations

To validate our theoretical results and compare our models to others in the literature, we performed a simulation study. Using Cellcoal [21] version 1.2.0, we generated two sets of 100 phylogenetic trees on 100 taxa with phased maternal and paternal DNA sequences; the first set with sequence length 1,000 and the second with sequence length 10,000. On each tree, maternal and paternal sequences evolved independently under the same GTR4 model, starting from a maternal and paternal sequence at the root which were each sampled independently from the stationary distribution of the GTR4 model. Note that this is not the default behavior in Cellcoal; by default, Cellcoal initializes the maternal sequence from the stationary distribution and copies this sequence to the paternal sequence.

Each of these simulations resulted in both 1) a true tree with branch lengths and 2) a collection of taxa with phased maternal-paternal sequences that have been simulated over that tree under independent maternal and paternal processes as described in Section 4.1. As such, the joint process is the Kronecker sum *Q*_ph_ = *Q*_*m*_ ⊕ *Q*_*p*_ and, as described in Section 4.2, since the maternal and paternal processes are equal, *Q*_ph_ can be lumped to form an unphased model *Q*_u_. Further, the generated phased sequences can also be considered as being generated under the unphased model by simply forgetting the phase information.

Using our implementation [22] of Felsenstein’s pruning algorithm [20], we used these generated sequences to find maximum likelihood estimates of rate parameters and branch lengths for the unphased lumped model and for the GT10 model from [12], under the assumption that the topology of the true tree is known. As a comparison, we also performed the same estimation using the RAXML-NG [23] software that forms an important component of Cellphy [12]. The results appear in Figures 3 and 4. In Figure 3a, we show the per-simulation differences between the true rate matrix and the estimated rate matrices for sequences of length 1K and 10K, while in Figure 3b we show mean of these differences for the 10K sequences. For the unphased lumped model, these simulations show the expected behavior. That is, there are moderate errors for the 1K sequences which decrease toward zero for the longer sequences. However, for the GT10_ML_ and GT10_Cellphy_ we see a different pattern. For these models, there is less variation in the estimates for longer sequences (Figure 3a) but the errors in the rate estimates do not tend toward zero and instead exhibit a structured bias in which the transitions between homozygous and heterozygous states (in both directions, see the north-east and south-west blocks shown in Figure 3b) are underestimated and the transitions between heterozygous states are overestimated (see the south-east blocks show in Figure 3b).

**Fig. 4:**
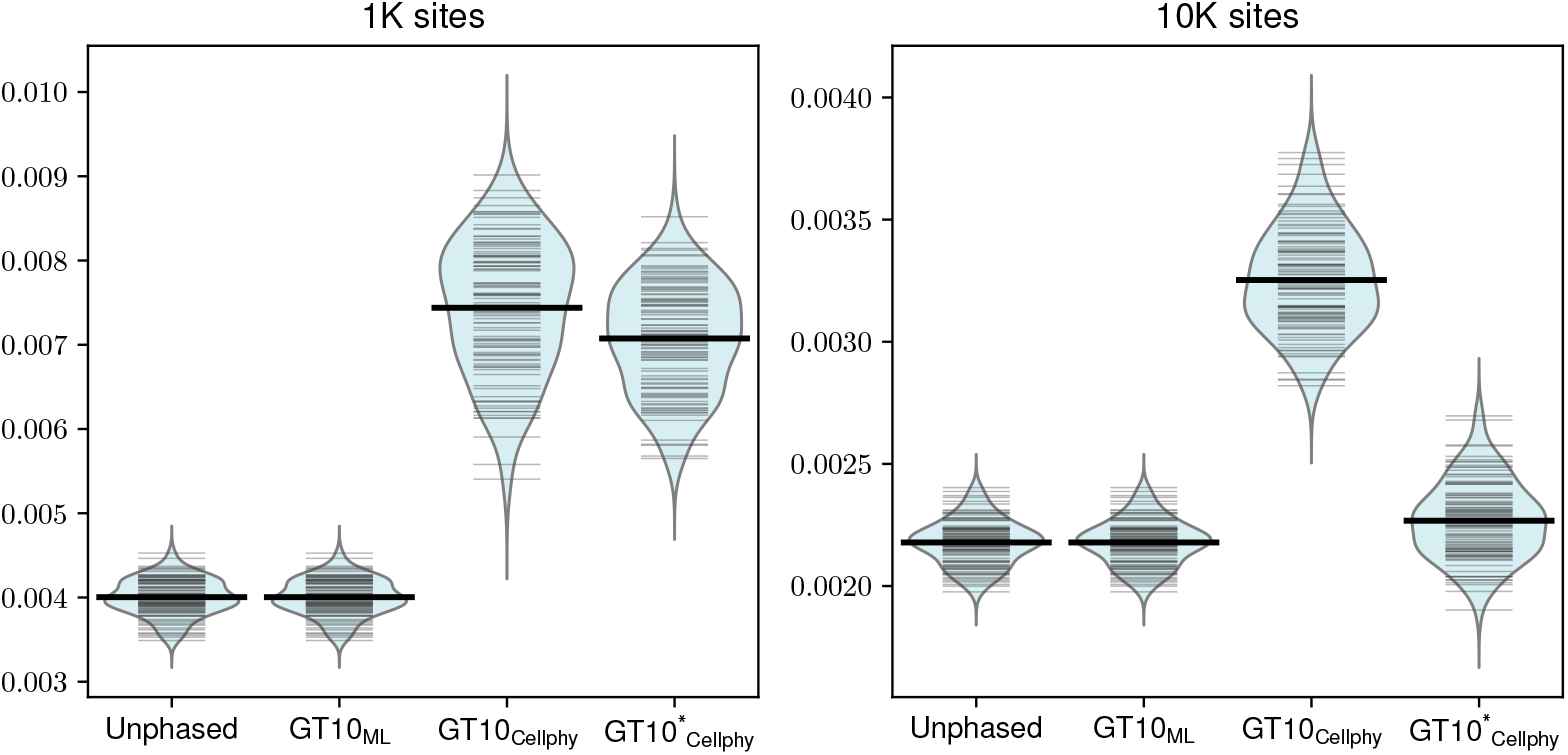
KF distance between inferred trees and the true tree. Here, the underlying topology of the trees are fixed at the true topology and so distances solely reflect the accuracy of branch-length estimation. The unphased lumped and GT10_ML_ models are estimated using our software [22] while GT10_Cellphy_ is estimated using RAXML-NG [12, 23]. The scaled GT10^*^_Cellphy_ estimate rescales the GT_Cellphy_ estimates to have an overall mutation rate compatible with the generative model, rather than the *µ* = 1 normalization for the GT10 model.

We measured the accuracy of branch length estimation by the Kuhner-Felsenstein (KF) distance [24], with the results show in 4. In addition to the estimates from the unphased lumped model, the GT10_ML_, and GT10_Cellphy_ model, we also display a scaled version of the GT10_Cellphy_ model. This scaled model is identical to the GT10_Cellphy_ model except for a scaling factor correction. This correction factor is necessary as RAXML-NG reports branch lengths scaled to be appropriate to a *µ* = 1 GT10 model. (See Section 4.6.) That is, these branch lengths are reported in a timescale that expects one mutation per unit time in a 10 state model, while the data was generated under the assumption of two independent processes with timescale that expect one mutation per unit time in each of their 4 state models. Further note that our software provides better branch length estimates in all cases. We are continuing to investigate why the unphased lumped model and GT10_ML_ models perform so similarly for branch lengths, despite the unphased lumped model giving a much better estimate of the true rate matrix.

The code for our simulations can be found at https://github.com/knappa/pruning.

## 9 Conclusions

While continuous time Markov models for haploid data are well studied, the general theory for diploid data is less well developed. Moreover, diploid data have several features that are not present in the haploid case such as phasing and marginalization to the maternal or paternal component processes. In this article, we primarily focused on the case where the diploid data were generated from independent maternal and paternal processes and were able to derive some useful results:

- If an unphased model is derived from lumping an independent maternal and paternal processes, then these processes must be identical. (Theorem 4.2)
- Theorem 4.2 gives an explicit formula for the rate matrix in terms of the lumping matrices. We present a ternary example in Equation 31.
- Equation 34 gives an explicit formula for the stationary distribution of the unphased lumped model in terms of the stationary distribution for the haploid models.
- Equation 41 expresses the mutation rate for the independent phased maternal-paternal process in terms of the mutation rates for the maternal and paternal processes.
- Equation 42 expresses the mutation rate for corresponding the unphased lumped process in terms of the rate for the maternal-paternal process.

Theorem 4.5 shows that a lumped process is time-reversible as long as the underlying process is time-reversible. Further, Equation 39 gives an explicit formula for the symmetric matrix representation for the independent maternal-paternal process, and Equation 40 gives an explicit formula for the unphased lumped process. As part of Section 7, we use these results to construct a symmetric matrix representation for our unphased lumped DNA model. This symmetric matrix naturally contains terms involving the rates of the maternal-paternal process and the nucleotide frequencies, and we compare it with the 10 state model in [12].

We also showed that, for the independent maternal-paternal phased process, the full likelihood function factors into a maternal and paternal likelihood. (Equation 46) This factorization does not hold for the unphased lumped process but, instead, the unphased likelihood is equivalent to a weighted sum of phased likelihoods, summed over all possible phase assignments. (Equation 48)

In Section 6, we gave examples of several alternative models on phased diploid data and explored their potential as unphased lumped models and their marginal lumpability properties. Similarly, in Section 7, we presented several unphased diploid models from the literature and evaluated the possibility of representing them as lumpings of an independent phased maternal-paternal process.

In Section 8, we presented numerical results comparing both our unphased lumped model and the GT10 model in [12]. These results demonstrate a clear improvement in rate parameter inference for the unphased lumped model. For branch length reconstruction, the results are more complicated. The unphased lumped model and GT10 performed similarly when inferred with our software. However, when inference was performed on the same dataset with RAXML-NG, a scaling subtlety was uncovered.

RAXML-NG uses a rate matrix normalization that fixes the overall *diploid* mutation rate to *µ* = 1. However, the simulated branch lengths were obtained under a generative model with a *haploid* mutation rate of *µ* = 1, which by Equation 42 corresponds to a *µ* = 2 diploid rate. We found that the branch-length reconstruction quality of RAXML-NG could be improved by taking this scaling into account.

This article raises several questions that we will address in future work, such as a characterization of dependent processes that are either maternal-paternal marginalizable, unphased lumpable, or both. We have also not addressed models that violate the site independence assumption. e.g. more realistic deletion models.

## Supporting information

Code for simulations

## Acknowledgments

This work was supported in part by the National Science Foundation under award No. DMS-2325775 (J.C., A.K.); and by the National Science Foundation under Grant No.

DMS-1929284 while the author Chifman was in residence at the Institute for Computational and Experimental Research in Mathematics in Providence, RI, during the Theory, Methods, and Applications of Quantitative Phylogenomics semester program (J.C.).

## Appendix A Select Proofs

### Lemma A.1

(Lemma 4.7). *Let X*(*t*) *be a continuous-time Markov process on a finite state space S with generator matrix Q and unique stationary distribution π. Suppose X*(*t*) *is lumpable with respect to a partition E of S with r classes. Then*,

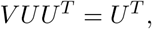

1. *D*_*π*_*V U* = *V UD*_*π*_, *where D*_*π*_ = diag(*π*), *and*
2. 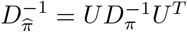, *where* 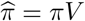 *and* 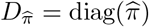.

*Proof* Recall from Equation 6 that *U* = (*V* ^*T*^ *V*)^−1^*V* ^*T*^ is a left inverse of *V*. Substituting for *U*^*T*^ and using *UV* = *I*_*r*_, we get

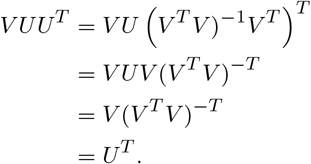

Now, suppose that for every partition class *E*_*k*_ ∈ *E* we have *π*_*i*_ = *π*_*j*_ for all *i, j* ∈ *E*_*k*_. Arrange the states of *S* according to the classes *E*_1_, *E*_2_, …, *E*_*r*_, and let *n*_*k*_ = |*E*_*k*_ |. Under this convention, we can write

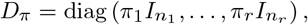

where *π*_*k*_ denotes the common value of *π*_*i*_ for *i* ∈ *E*_*k*_. The matrix *V U* is also block diagonal, with the block corresponding to class *E*_*k*_ given by 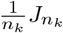, where *J*_*n*_ is the all-ones matrix. In particular,

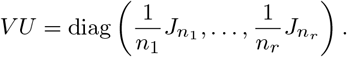

Direct computation now shows property (1):

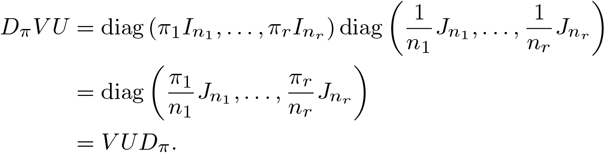

Part (2) now follows from part (1) and the identity 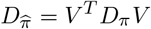:

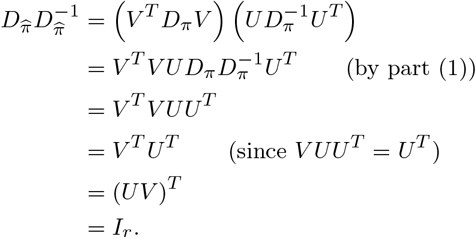

Thus we conclude that the inverse of 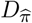 is 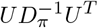. □

### Identities A.2.

*Let V be the matrix associated with the unphased partition G*_u_, *as defined in Section 4.2. Define the Kronecker product of two standard column vectors e*_*i*_, *e*_*j*_ ∈ ℝ ^*κ*^ *by e*_*ij*_ := *e*_*i*_ ⊗ *e*_*j*_. *Then*

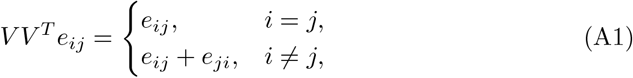

*Proof* Unphased partition *G*_u_ has size 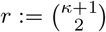, consisting of *κ* classes of the form *i/i* and 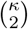 classes of the form *i/j*. The corresponding matrix *V* has size *κ*^2^ *× r*, with rows indexed by phased states and columns indexed by unphased classes. The rows of *V* are standard vectors in ℝ ^*r*^, with repeated rows corresponding to phased states *i*_m_*j*_p_ and *j*_m_*i*_p_, where *i* ≠*j*. Assume the phased states are ordered lexicographically, so that *e*_*ij*_ = *e*_*i*_ ⊗ *e*_*j*_ is the standard vector with 1 in position (*i* − 1)*κ* + *j*.

Since multiplication of a matrix by a standard vector selects the corresponding column, *V* ^*T*^ *e*_*ij*_ is the row of *V* indexed by the phased state *i*_m_*j*_p_. Consequently, *V V* ^*T*^ *e*_*ij*_ is the column of *V* corresponding to the unphased class containing *i*_m_*j*_p_.

If *i* = *j*, this class is *i/i*, which contains only the phased state *i*_m_*i*_p_. Hence,

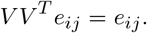

If *i* ≠ *j*, this class is *i/j*, which contains the two phased states *i*_m_*j*_p_ and *j*_m_*i*_p_. Therefore,

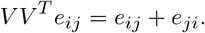

□

### Identities A.3.

*Let V be an n × r matrix such that each row of V is a standard vector in* ℝ ^*r*^. *Let A and B be matrices of size r × m. Then*

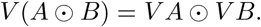

*Proof* Consider a single row of *V*. Since each row of *V* is a standard vector, left multiplication by that row selects the corresponding row of a matrix. Since the Hadamard product is defined entrywise, selecting a row of *A*⊙ *B* is the same as selecting the corresponding rows of *A* and *B* and then taking their Hadamard product. □

## Appendix B Site Pattern Probability using the Pointwise Product

Consider a rooted three-taxon tree *T* as depicted in Figure B1. Let *X*_ℓ_ denote the state observed for lineage ℓ = 1, 2, 3, and let *P* (*t*_*i*_) denote the transition probability matrix associated with edge *t*_*i*_. There are multiple equivalent ways to compute the joint probability (site pattern probability)

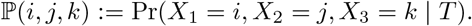

As explained in Section 5, we use Felsenstein’s pruning algorithm [20] to compute site pattern probabilities, but with a slight difference in notation, since for our models it is more convenient to express them via the pointwise Hadamard product of matrices. In what follows, we do not try to prove the validity of the pruning algorithm, but merely to justify our notation.

**Fig. B1.**
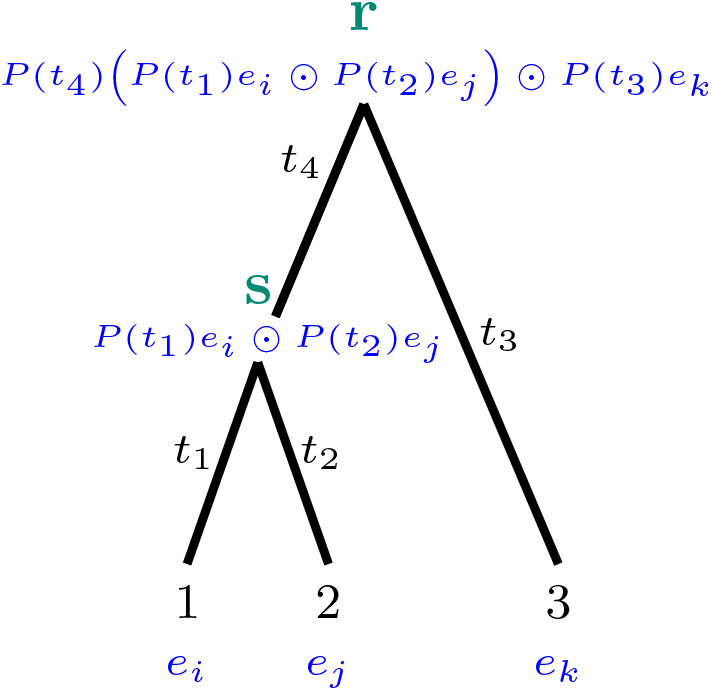
Rooted 3-taxon tree.

Consider a continuous-time finite-state Markov model of sequence evolution with state space *S* = {1, 2, …, *n}*, and let the standard basis column vector *e*_*i*_ ∈ ℝ^*n*^ correspond to state *i* ∈ *S*. Thus, for a site pattern *X*_1_*X*_2_*X*_3_ = *ijk* at the tips of the tree, we assign the standard vectors *e*_*i*_, *e*_*j*_, and *e*_*k*_.

To each vertex *v* in *T*, Felsenstein’s algorithm assigns an *n*-dimensional vector **v** of conditional probabilities of the observed data on descendants of *v*, conditional on the state at *v* and the tree. For example, on *s*, the components of **s** are

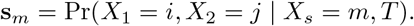

Similarly, for *r*, the components of **r** are

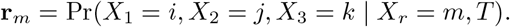

This description is compatible with the previous statements about the vectors for the tips as long as we consider a tip its own descendant. The appearance of the Hadamard product can now be seen as a result of independent evolution on the subtrees hanging off of *v*. That is, for *s*, independent evolution along the edges leading to tips 1 and 2 means that

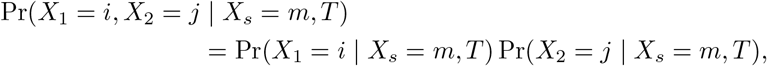

which we immediately recognize as the Hadamard product written component-wise. That is, if we write ⊙ for the Hadamard product,

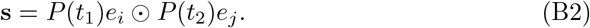

Similarly, for *r*, independent evolution gives us

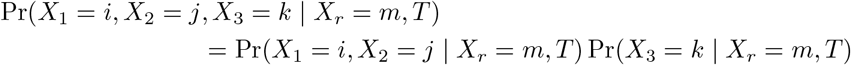

which again can be expressed as a Hadamard product of vectors. So, with **s** from Equation B2 and the vector *e*_*k*_ corresponding to tip 3 with transition probability matrices *P* (*t*_4_) and *P* (*t*_3_) respectively, we see that

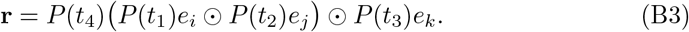

**r** = *P* (*t*_4_) *P* (*t*_1_)*e*_*i*_ ⊙ *P* (*t*_2_)*e*_*j*_ ⊙ *P* (*t*_3_)*e*_*k*_. (B3) Finally, the probability of observing the site pattern *ijk* at the tips of the tree is:

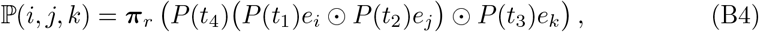

where ***π***_*r*_ is the probability distribution of states at the root. One can see that this can be extended to any binary tree, with more factors appearing as one climbs up the tree.

## Appendix C 16 state models lumping to 10 state Cellphy model

Let *Q*_*GT* 10_ be the rate matrix for a particular parameterization of the 10 state Cellphy model [12] as discussed in Section 7. Every unphased-lumpable diploid DNA model which lumps to *Q*_*GT* 10_ is of the form *Q*_∗_ +*R* described below. There are 12 free parameters *r*_1_, …, *r*_12_ whose only constraint is that the rates must remain non-negative. *e*.*g*. 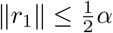. The first matrix, *Q*_∗_, is of the form *MD* where

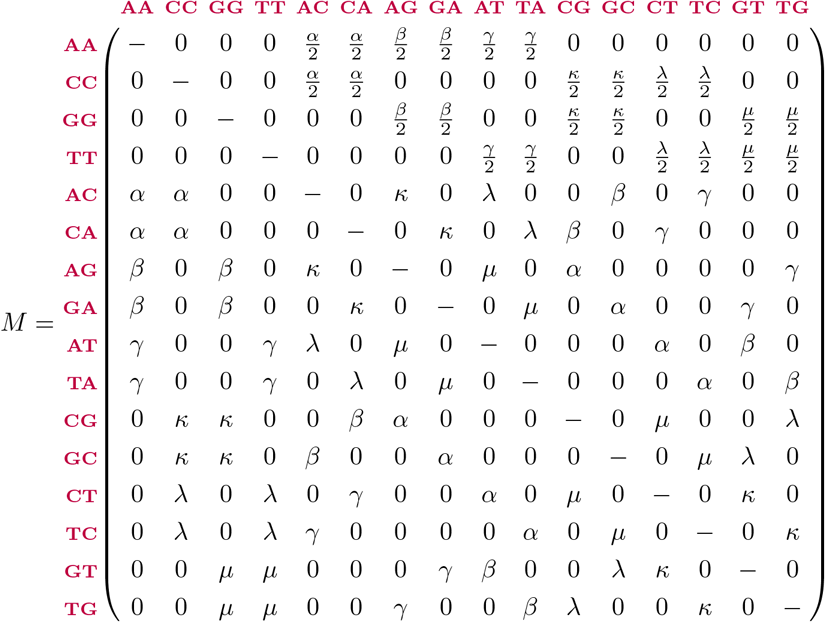

and *D* is the diagonal matrix with diagonal entries *π*_*aa*_, *π*_*cc*_, *π*_*gg*_, *π*_*tt*_, *π*_*ac*_, *π*_*ca*_, *π*_*ag*_, *π*_*ga*_, *π*_*at*_, *π*_*ta*_, *π*_*cg*_, *π*_*gc*_, *π*_*ct*_, *π*_*tc*_, *π*_*gt*_, and *π*_*tg*_, matching the column states. This distribution must be phase-invariant. *e*.*g. π*_*ac*_ = *π*_*ca*_. Although we have presented *Q*_∗_ similarly to the symmetric matrix repersentation for time-reversible rate matrices, *M* is not symmetric and *Q*_∗_ is not time-reversible.

The second matrix, *R*, contains the free variables and is a block matrix with nonzero entries only in the upper right quadrant. Similarly to *Q*_∗_, we can express *R* in the form *R* = *ND*. Let *D*^*′*^ be the diagonal matrix with entries *π*_*ac*_, *π*_*ca*_, *π*_*ag*_, *π*_*ga*_, *π*_*at*_, *π*_*ta*_, *π*_*cg*_, *π*_*gc*_, *π*_*ct*_, *π*_*tc*_, *π*_*gt*_, and *π*_*tg*_, matching the heterozygous column states. Then we then get

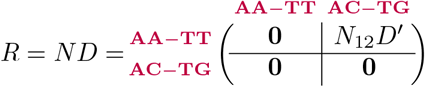

with

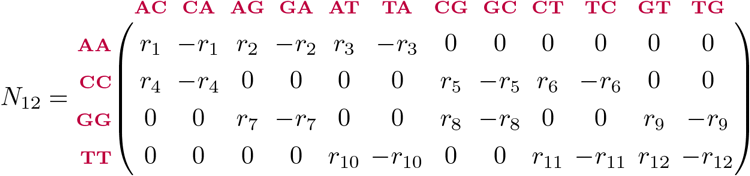

