## Supplementary figures and images for "Generalized Maternal-Paternal Genotype Markov Model for Single-Cell Lineages"

### dowload-link.png

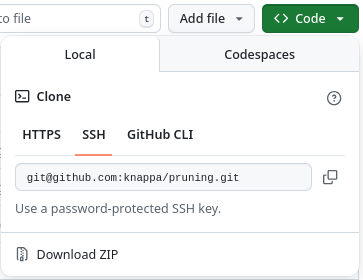

### model-comparisons-1K.pdf

Unphased DNA

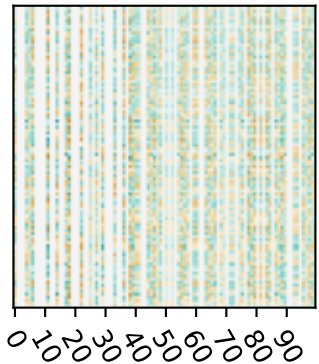

Cellphy

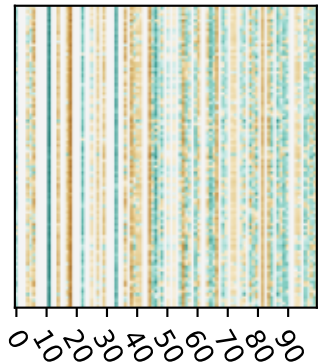

GTR10Z

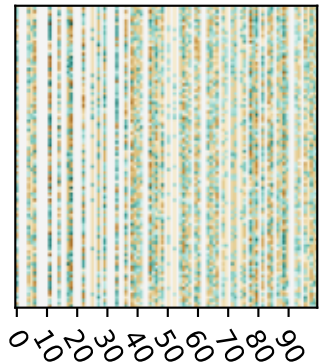

GTR10

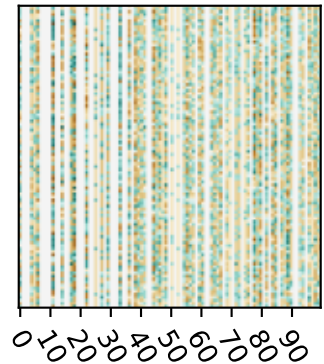

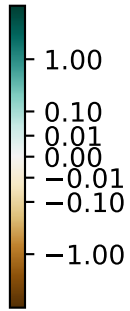

### model-comparisons-10K.pdf

Unphased DNA

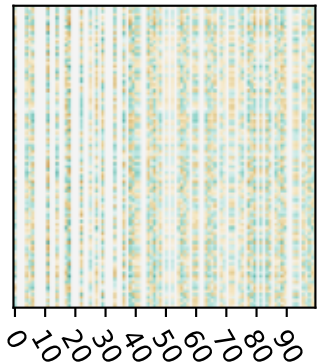

Cellphy

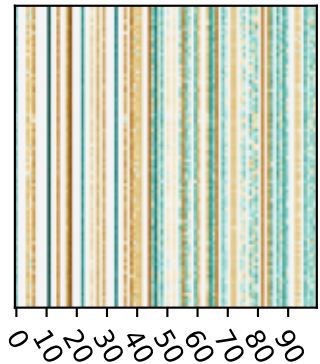

GTR10Z

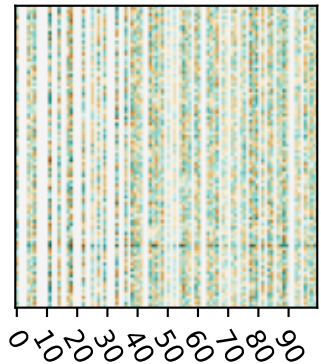

GTR10

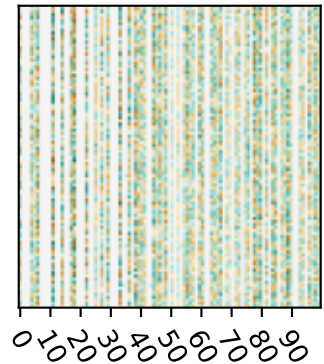

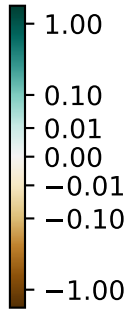
